# Immunogenicity and efficacy of the COVID-19 candidate vector vaccine MVA SARS 2 S in preclinical vaccination

**DOI:** 10.1101/2021.01.09.426032

**Authors:** Alina Tscherne, Jan Hendrik Schwarz, Cornelius Rohde, Alexandra Kupke, Georgia Kalodimou, Leonard Limpinsel, Nisreen M.A. Okba, Berislav Bošnjak, Inga Sandrock, Sandro Halwe, Lucie Sauerhering, Katrin Brosinski, Nan Liangliang, Elke Duell, Sylvia Jany, Astrid Freudenstein, Jörg Schmidt, Anke Werner, Michelle Gellhorn Sera, Michael Klüver, Wolfgang Guggemos, Michael Seilmaier, Clemens-Martin Wendtner, Reinhold Förster, Bart L. Haagmans, Stephan Becker, Gerd Sutter, Asisa Volz

**Affiliations:** Division of Virology, Department of Veterinary Sciences, LMU Munich, Munich, Germany; Institute of Virology, Philipps University Marburg, Marburg, Germany; German Center for Infection Research, Munich, Germany; German Center for Infection Research, Gießen-Marburg-Langen, Germany; Department of Viroscience, Erasmus Medical Center, Rotterdam, The Netherlands; Institute of Immunology, Hannover Medical School, Hannover, Germany; German Center for Infection Research, Hannover, Germany; Cluster of Excellence RESIST (EXC 2155), Hannover Medical School, Hannover, Germany; Munich Clinic Schwabing, Academic Teaching Hospital, LMU Munich, Munich, Germany; Institute of Virology, University of Veterinary Medicine Hannover, Germany

**Keywords:** vaccine vector, vaccinia virus, poxvirus, non-clinical testing

## Abstract

The severe acute respiratory syndrome (SARS) coronavirus 2 (SARS-CoV-2) has emerged as the infectious agent causing the pandemic coronavirus disease 2019 (COVID-19) with dramatic consequences for global human health and economics. Previously, we reached clinical evaluation with our vector vaccine based on vaccinia virus MVA against the Middle East respiratory syndrome coronavirus (MERS-CoV), which causes an infection in humans similar to SARS and COVID-19. Here, we describe the construction and preclinical characterization of a recombinant MVA expressing full-length SARS-CoV-2 spike (S) protein (MVA-SARS-2-S). Genetic stability and growth characteristics of MVA-SARS-2-S, plus its robust synthesis of S antigen, make it a suitable candidate vaccine for industrial scale production. Vaccinated mice produced S antigen-specific CD8+ T cells and serum antibodies binding to S glycoprotein that neutralized SARS-CoV-2. Prime-boost vaccination with MVA-SARS-2-S protected mice sensitized with a human ACE2-expressing adenovirus from SARS-CoV-2 infection. MVA-SARS-2-S is currently being investigated in a phase I clinical trial as aspirant for developing a safe and efficacious vaccine against COVID-19.

**Significance Statement:** The highly attenuated vaccinia virus MVA is licensed as smallpox vaccine, and as vector it is a component of the approved Adenovirus-MVA-based prime-boost vaccine against Ebola virus disease. Here we provide results from testing the COVID-19 candidate vaccine MVA-SARS-2-S, a poxvirus-based vector vaccine that proceeded to clinical evaluation. When administered by intramuscular inoculation, MVA-SARS-2-S expresses and safely delivers the full-length SARS-CoV-2 spike (S) protein, inducing balanced SARS-CoV-2-specific cellular and humoral immunity, and protective efficacy in vaccinated mice. Substantial clinical experience has already been gained with MVA vectors using homologous and heterologous prime-boost applications, including the immunization of children and immunocompromised individuals. Thus, MVA-SARS-2-S represents an important resource for developing further optimized COVID-19 vaccines.

## Introduction

The severe acute respiratory syndrome coronavirus 2 (SARS-CoV-2), the causal agent of coronavirus disease 2019 (COVID-19), first emerged in late 2019 in China (1). SARS-CoV-2 exhibits extremely efficient human-to-human transmission, the new pathogen rapidly spread worldwide, and within months caused a global pandemic, changing daily life for billions of people. The COVID-19 case fatality rate of ~2-5% makes the development of countermeasures a global priority. In fact, the development of COVID-19 vaccine candidates is advancing at an international level with unprecedented speed. About a year after the first known cases of COVID-19 we can account for >50 SARS-Cov-2-specific vaccines in clinical evaluations and >10 candidate vaccines already in phase III trials (2–4). Yet, we still lack information on the key immune mechanisms needed for protection against COVID-19. A better understanding of the types of immune response elicited upon natural SARS-CoV-2 infections has become an essential component to assess the promise of various vaccination strategies (5).

The SARS-CoV-2 spike (S) protein serves as the most important target antigen for vaccine development based on preclinical research on candidate vaccines against SARS-CoV or MERS-CoV. The trimeric S glycoprotein is a prominent structure at the virion surface and essential for SARS-CoV-2 cell entry. As a class I viral fusion protein, it mediates virus interaction with the cellular receptor angiotensin-converting enzyme 2 (ACE2), and fusion with the host cell membrane, both key steps in infection. Thus infection can be prevented by S-specific antibodies neutralizing the virus (6–9).

Among the front-runner vaccines are new technologies such as messenger RNA (mRNA)-based vaccines and non-replicating adenovirus vector vaccines (10). First reports from these SARS-CoV-2 S-specific vaccines in phase 1/2 clinical studies show acceptable safety and promising immunogenicity profiles, and by now first data from large phase 3 clinical trials show promising levels of protective efficacy (4, 11–13). This is good news because efficacious vaccines will provide a strategy to change SARS-CoV-2 transmission dynamics. In addition, multiple vaccine types will be advantageous to meet specific demands across different target populations. This includes the possibility of using heterologous immunization strategies depending on an individual’s health status, boosting capacities and the need for balanced humoral and Th1-directed cellular immune responses.

Modified Vaccinia virus Ankara (MVA), a highly attenuated strain of vaccinia virus originating from growth selection on chicken embryo tissue cultures, shows a characteristic replication defect in mammalian cells but allows unimpaired production of heterologous proteins (14). At present, MVA serves as an advanced vaccine technology platform for developing new vector vaccines against infectious disease and cancer including emerging viruses (15). In response to the ongoing pandemic, the MVA vector vaccine platform allows rapid generation of experimental SARS-CoV-2-specific vaccines (16). Previous work from our laboratory addressed the development of an MVA candidate vaccine against MERS with immunizations in animal models demonstrating the safety, immunogenicity and protective efficacy of MVA-induced MERS-CoV S-antigen specific immunity (17–20). Clinical safety and immunogenicity of the MVA-MERS-S candidate vaccine was established in a first-in-man phase I clinical study under funding from the German Center for Infection Research (DZIF) (21).

Here, we show that a recombinant MVA virus produces the full-length S protein of SARS CoV-2 as ~190-200 kDa N-glycosylated protein. Our studies confirmed cleavage of the mature full-length S glycoprotein into an amino-terminal domain (S1) and a ~80-100 kDa carboxy-terminal domain (S2) that is anchored to the membrane. When tested as a vaccine in BALB/c mice, recombinant MVA expressing the S glycoprotein induced SARS-CoV-2-specific T cells and antibodies, and robustly protected vaccinated animals against lung infection upon SARS-CoV-2 challenge.

## Results

### Design and generation of candidate MVA vector viruses

cDNA containing the entire gene sequence encoding SARS-CoV-2 S (SARS-2-S) from the virus isolate Wuhan HU-1 (GenBank accession no. MN908947.1) was placed under the transcriptional control of the enhanced synthetic vaccinia virus early/late promoter PmH5 (22) in the MVA vector plasmid pIIIH5red-SARS-2-S, and introduced by homologous recombination into deletion site III in the MVA genome (Fig. 1A). Clonal recombinant MVA viruses expressing SARS-2-S (MVA-SARS-2-S) were isolated in repetitive plaque purification using transient coproduction of the fluorescent marker protein mCherry to screen for red fluorescent cell foci (17, 23). PCR analysis of viral DNA confirmed the genetic integrity of the recombinant viruses demonstrating the site-specific insertion of the heterologous SARS-2-S gene sequences in the MVA genome, and subsequently the proper removal of the mCherry marker gene from the genome of final recombinant viruses (Fig. 1*B*). MVA-SARS-2-S virus isolates were genetically stable and showed the expected MVA–specific genetics with regard to characteristic deletions and sequence alterations in the MVA genome (SI *Appendix*, Fig. S1). The recombinant viruses replicated efficiently in the chicken embryo fibroblast cell line DF-1, but not in the human cell lines HeLa, A549 or HaCat (Fig. 1*C*).

**Fig. 1.**
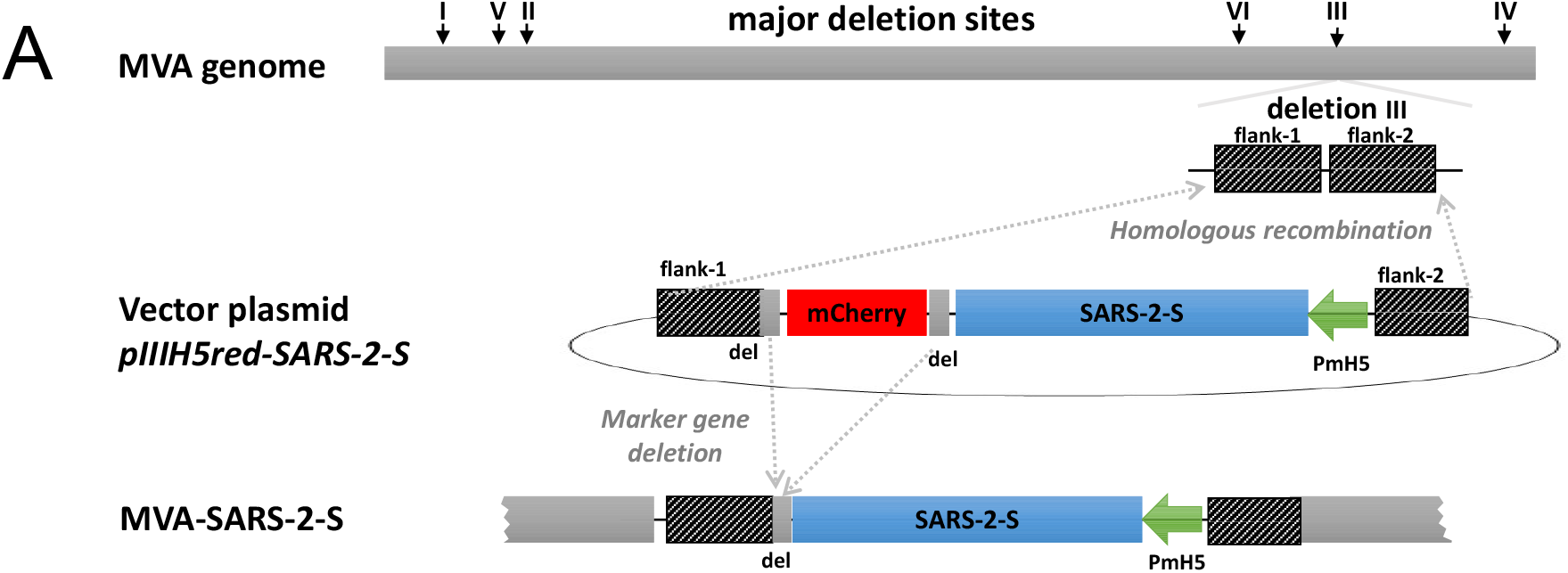
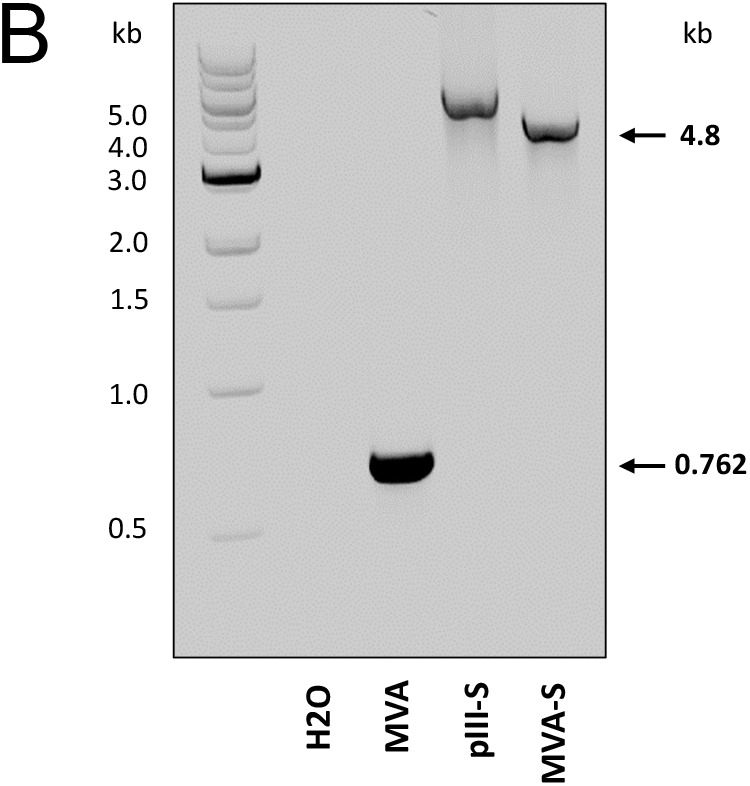
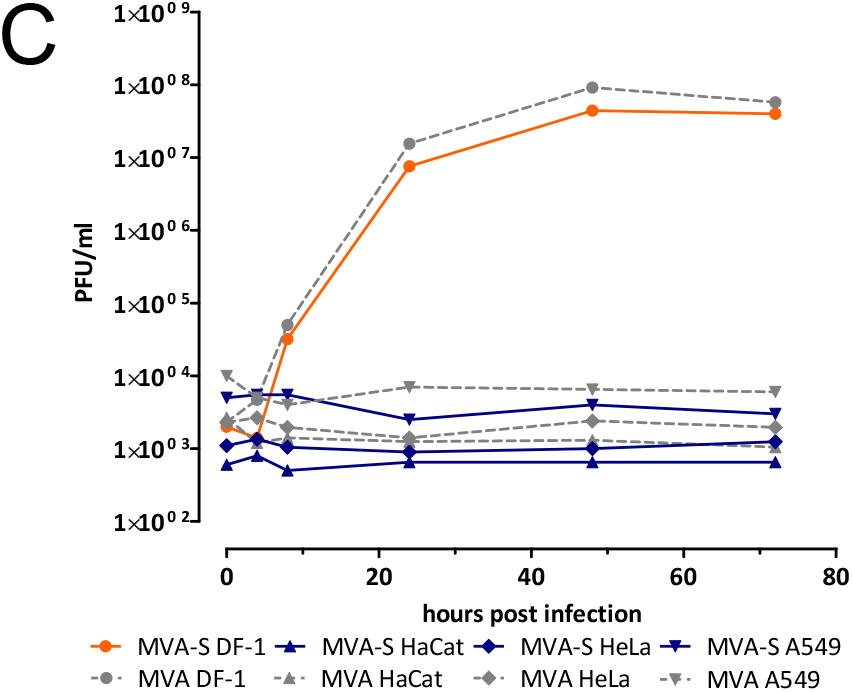
Construction and virological characterization of MVA-SARS-2-S. **(A)** Schematic diagram of the MVA genome with the major deletion sites I to VI. The site of deletion III was targeted for insertion of the gene sequence encoding the S protein of SARS-CoV-2 isolate Wuhan-HU-1 (SARS-2-S). SARS-2-S was placed under transcriptional control of the vaccinia virus promoter PmH5 within the MVA vector plasmid pIIIH5red-SARS-2-S. Insertion occurred via homologous recombination between MVA DNA sequences (flank-1 and flank-2) adjacent to deletion site III in the MVA genome and copies cloned in the vector plasmid. MVA-SARS-2-S was isolated by plaque purification screening for co-production of the red fluorescent marker protein mCherry. A repetition of short flank-1 derived DNA sequences (del) served to remove the marker gene by intragenomic homologous recombination (marker gene deletion). **(B)** Genetic integrity of MVA-SARS-2-S (MVA-S). PCR analysis of genomic viral DNA confirmed stable insertion of the SARS-2-S sequence into deletion site III of the MVA genome. The precise intragenomic deletion of the marker gene mCherry during plaque purification was revealed by amplification of a PCR product with the expected molecular weight (4.8 kb) from MVA-S genomic DNA compared to the large product amplified from pIIIH5red-SARS-2-S plasmid DNA template (pIII-S). The deletion III sitespecific oligonucleotide primers amplified a characteristic 0.762 kb DNA fragment from genomic, non-recombinant MVA DNA. **(C)** Multiple-step growth analysis of recombinant MVA-SARS-2-S (MVA-S) and non-recombinant MVA (MVA). Cells were infected at a multiplicity of infection (MOI) of 0.05 with MVA-S or MVA and collected at the indicated time points. Titration was performed on CEF cells and plaque-forming units (PFU) were determined. MVA-S and MVA could be efficiently amplified on DF-1 but failed to productively grow on cells of human origin (HaCat, HeLa and A549).

### Characterization of SARS-CoV-2 S expressed by recombinant MVA

To determine the expression pattern of the recombinant SARS-CoV-2 S protein, we stained MVA-SARS-2-S infected Vero cells with HAtag or S-specific monoclonal antibodies and analyzed them using fluorescence microscopy. A mouse monoclonal antibody directed against the nine amino acid-HA-tag at the carboxyl (C)-terminus of the recombinant SARS-2-S protein revealed highly specific staining in permeabilized cells corresponding to the expected intracellular localization of the S protein C-terminal end. A SARS-CoV-1 S-specific monoclonal antibody recognizing an epitope in the external domain of the S protein also allowed the specific staining of non-permeabilized MVA-SARS-2-S infected Vero cells, suggesting that the SARS-2-S protein was readily translocated to the cytoplasma membrane (Fig. 2*A*).

**Fig. 2.**
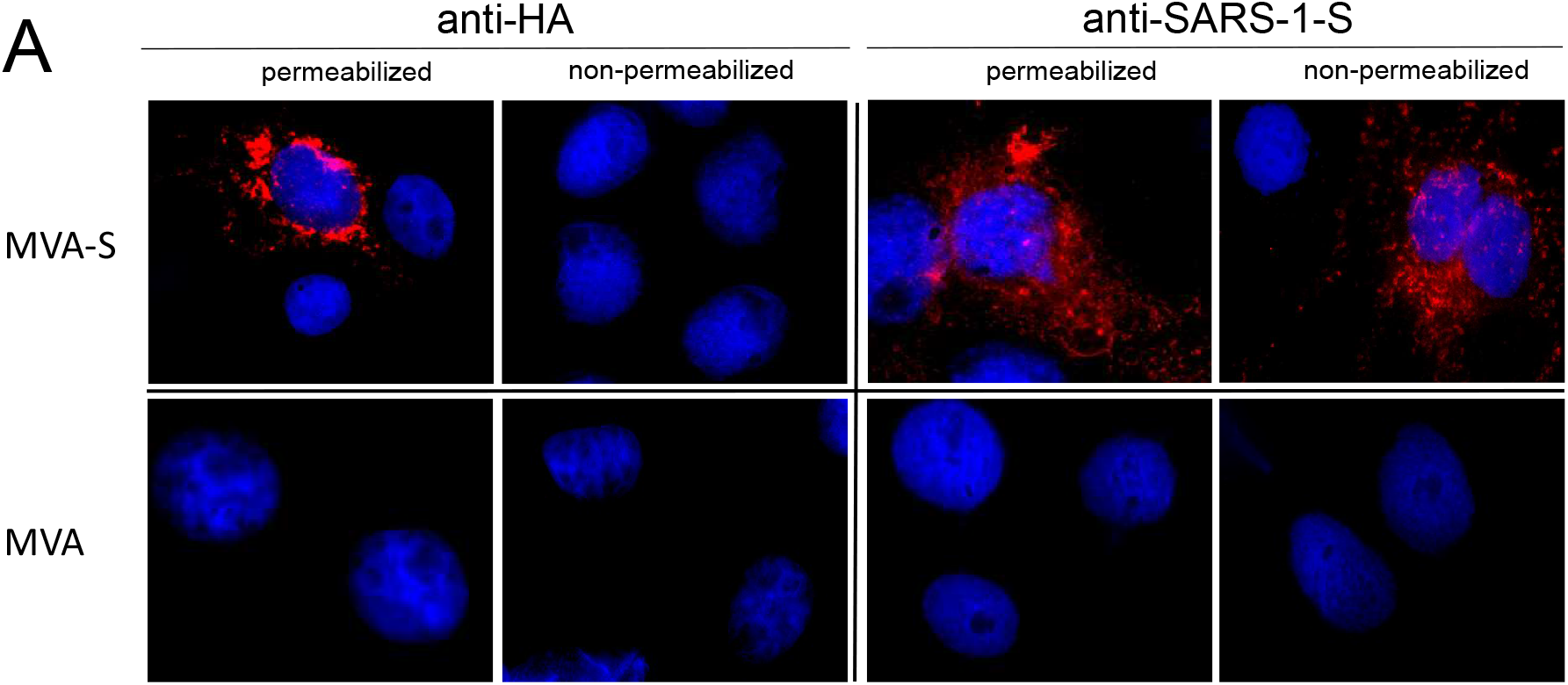
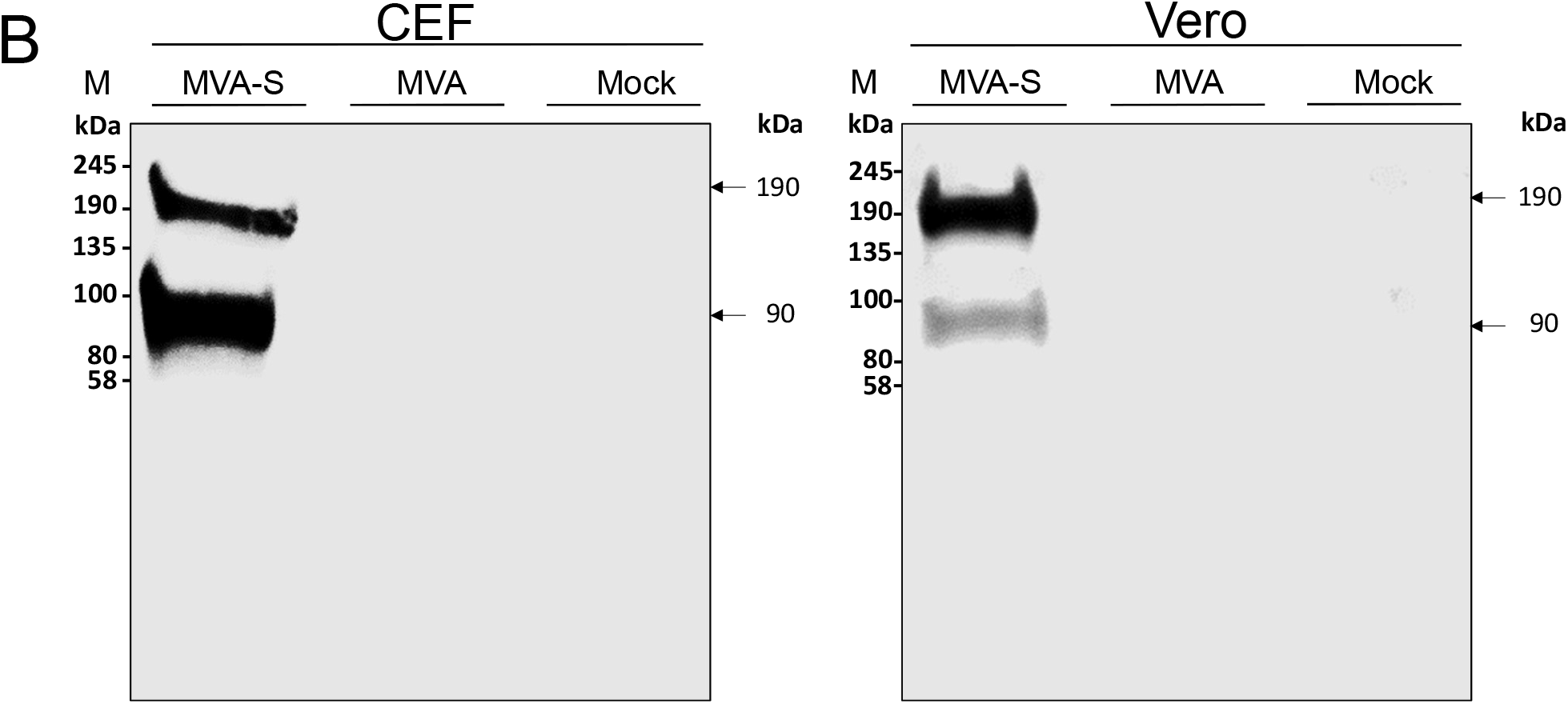
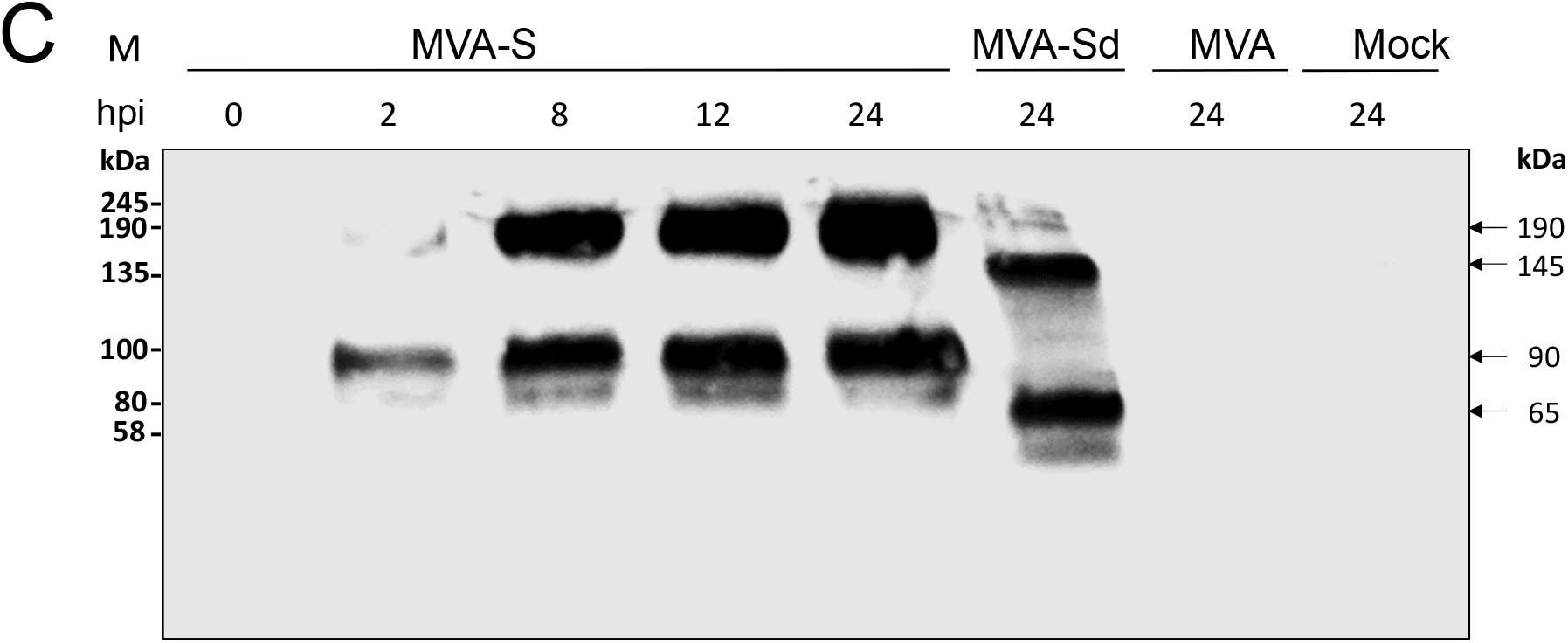
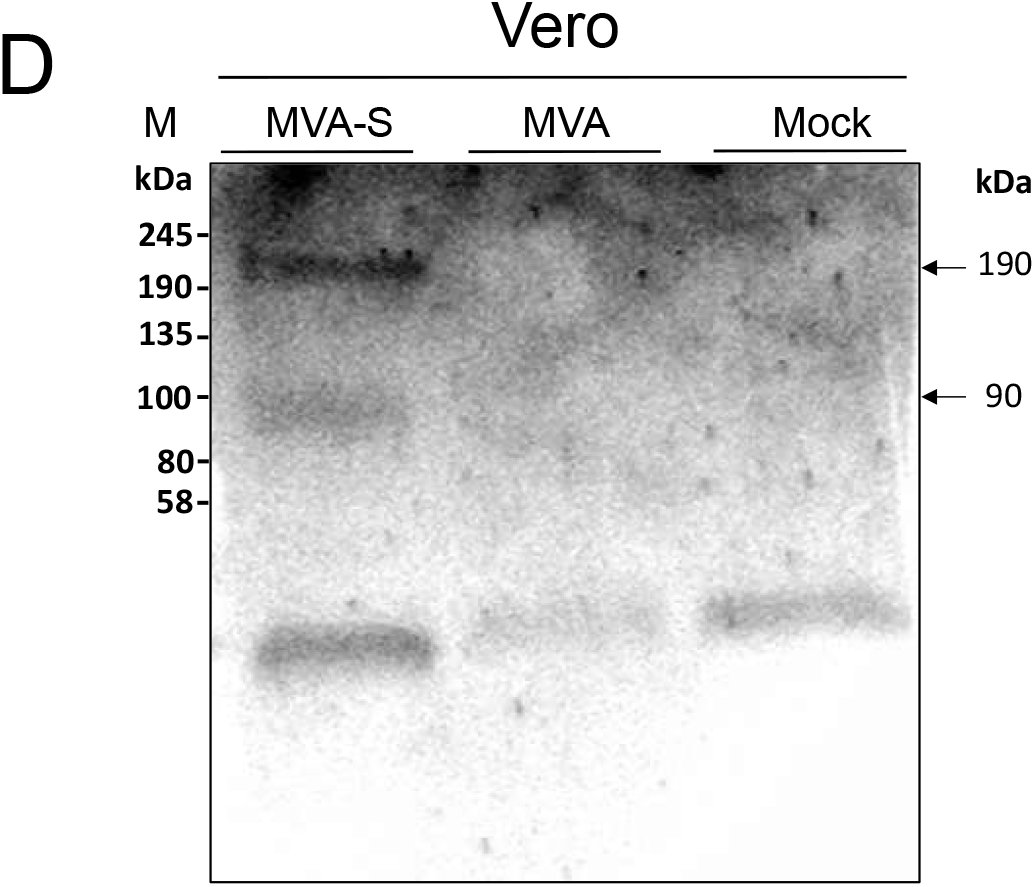
Synthesis of full-length Spike glycoprotein in MVA-SARS-2-S (MVA-S) infected cells. **(A)** Cells were infected at a multiplicity of infection of 0.5. MVA infected cells served as controls. Paraformaldehyde fixed cells were either permeabilized or non-permeabilized and probed with mouse monoclonal antibodies directed against the HAtag or the S protein of SARS-Cov-1 (SARS-1-S). Polyclonal goat anti-mouse secondary antibody was used for S-specific fluorescent staining (red). Cell nuclei were counterstained with DAPI (blue). **(B)** Chicken embryonic fibroblasts (CEF) and Vero cells were infected with a multiplicity of infection (MOI) of 10 and collected 24 hours post infection (hpi). **(C)** Vero cells were infected with MVA-SARS-2-S (MVA-S) at a MOI of 10 and collected after the indicated time points. Deglycosylation with PNGase F (MVA-Sd) was performed with the sample collected after 24 hpi. Polypeptides in cell lysates were separated by SDS-PAGE and analyzed with a monoclonal antibody against the HA-tag (1:8000) (a, b) or with human serum (1:200) **(D)** Lysates from non-infected (Mock) or non-recombinant MVA infected (MVA) cells were used as controls.

To examine the MVA-produced recombinant S glycoprotein in more detail, we prepared total lysates from MVA-SARS-2-S infected CEF or Vero cells for separation by SDS-PAGE and subsequent immunoblot analysis (Fig. 2). The mouse monoclonal antibody directed against the HAtag at the C-terminus of the recombinant SARS-2-S protein revealed two prominent protein bands that migrated with molecular masses of approximately 190 kDa and 90-100 kDa (Fig 2*B*). As in the SDS-PAGE the detected protein bands migrated at molecular masses significantly higher than the 145 kDa predicted for full-length SARS-CoV-2 S protein based on its amino acid sequence, we hypothesized that the proteins might be glycosylated. Indeed, NetNGlyc 1.0 server analysis indicated the presence of at least 17 N-glycosylation sites for co-and post-translational modifications. The treatment of cell lysates with peptide-N-glycosidase F (PNGase F), which removes all N-linked oligosaccharide chains from glycoproteins, reduced the molecular masses of the recombinant SARS-2-S protein bands from 190 kDa to 145 kDa and from 90-100 kDa to 65 kDa, matching the expected sizes of unmodified SARS-CoV-2 S and the S2 cleavage product, respectively (Fig 2*C*).

Interestingly, the S2 cleavage band was more prominent in the lysates from infected CEF cells, whereas lysates from infected Vero cells contained more full-length protein, suggesting host cell-specific differences in the proteolytic cleavage of the S protein (Fig. 2*B*). More importantly, both isoforms were detectable as early as 2 hours post-infection (hpi), indicating proper early transcription from the synthetic MVA promoter PmH5, and their amount increased up to 24 hpi, consistent with the timing of abundant vaccinia viral late protein synthesis (Fig. 2*C*). Moreover, Western blot analysis using serum antibodies from a COVID-19 patient hospitalized with pneumonia also revealed protein bands corresponding to the molecular masses of full-length S protein or the S2 polypeptide (Fig. 2*D*).

### MVA-SARS-2-S induced antibody responses in mice

To evaluate whether MVA-SARS-CoV-2 induces SARS-CoV-2 specific antibodies, we vaccinated BALB/c mice with a low-dose (LD) or high-dose (HD) of MVA-SARS-CoV-2 (10^7^ or 10^8^ plaque-forming units (PFU), respectively) using intramuscular (i.m.) administration and prime-boost immunization schedules with a 3-week interval (Fig. 3, *SI Appendix*, Fig. S2). At day 18 after the prime inoculation, we detected serum IgG antibodies binding to whole recombinant SARS-CoV-2 S protein in the sera from 3/8 LD-vaccinated and 4/6 HD-vaccinated animals by ELISA (Fig. 3*A*). Following the booster immunization on day 21, all vaccinated animals mounted high levels of S-binding serum IgG antibodies with mean titers of 1:900 for the LD vaccination group and 1:1257 for the HD group (Fig. 3*A*). More importantly, sera from vaccinated mice also contained antibodies binding to the S protein receptor-binding domain (RBD). Already at day 18 post priming, the RBD-binding antibodies were detected in 33% of the mice in the LD dose group (2/6 mice, mean OD value 0.35) and 50% of the mice receiving the HD immunization (3/6, mean OD 0.63). The boost vaccinations increased the levels of RBD specific antibodies with 87.5% seropositive mice in the 10^7^ dose group (7/8, mean OD 1.81) and 100% of the animals vaccinated with 10^8^ PFU MVA-SARS2-S (8/8, mean OD 2.92) (Figure 3*B*). Since live virus neutralization is the gold standard for coronavirus serologic analysis, we next assessed the mouse sera in two different assays for SARS-CoV-2 neutralization, a plaque reduction neutralization test 50 (PRNT_50_) (24) and a complete virus neutralization test (VNT_100_) (9) (Fig. 3 *C* and *D*). On day 18 following prime immunization the PRNT_50_ revealed low amounts of SARS-CoV-2 neutralizing antibodies in 50-80% of the sera from vaccinated animals (PRNT_50_ titres of 20-40 for both dose groups). After the boost vaccinations we detected neutralizing activities in all sera from MVA-SARS-2-S vaccinated mice with average PRNT_50_ titres of 117 (LD) and 600 (HD) (Fig. 3*C*). Using the VNT_100_ assay we detected neutralizing activities in 79% of all sera following MVA-SARS-2-S booster immunizations with mean reciprocal titres of 19.8 (4/6 seropositive mice, LD group) and 105.8 (7/8 mice, HD group) (Fig. 3*D*). We obtained similar results when testing the sera in a recently established high-throughput surrogate virus neutralization test for SARS CoV-2 (sVNT)(25). After the boost immunizations on day 21, we detected levels of surrogate neutralizing antibodies with mean titers of 400 (4/6 seropositive mice, LD) and 840 (6/6, HD) (Fig. 3*E*, *SI Appendix*, Fig. S3). Altogether, these results indicate that both LD and HD 3-week prime-boost immunization protocols induce a robust anti-SARS-CoV2-S humoral response and lead to generation of neutralizing anti-SARS-CoV2-S antibodies.

**Fig. 3.**
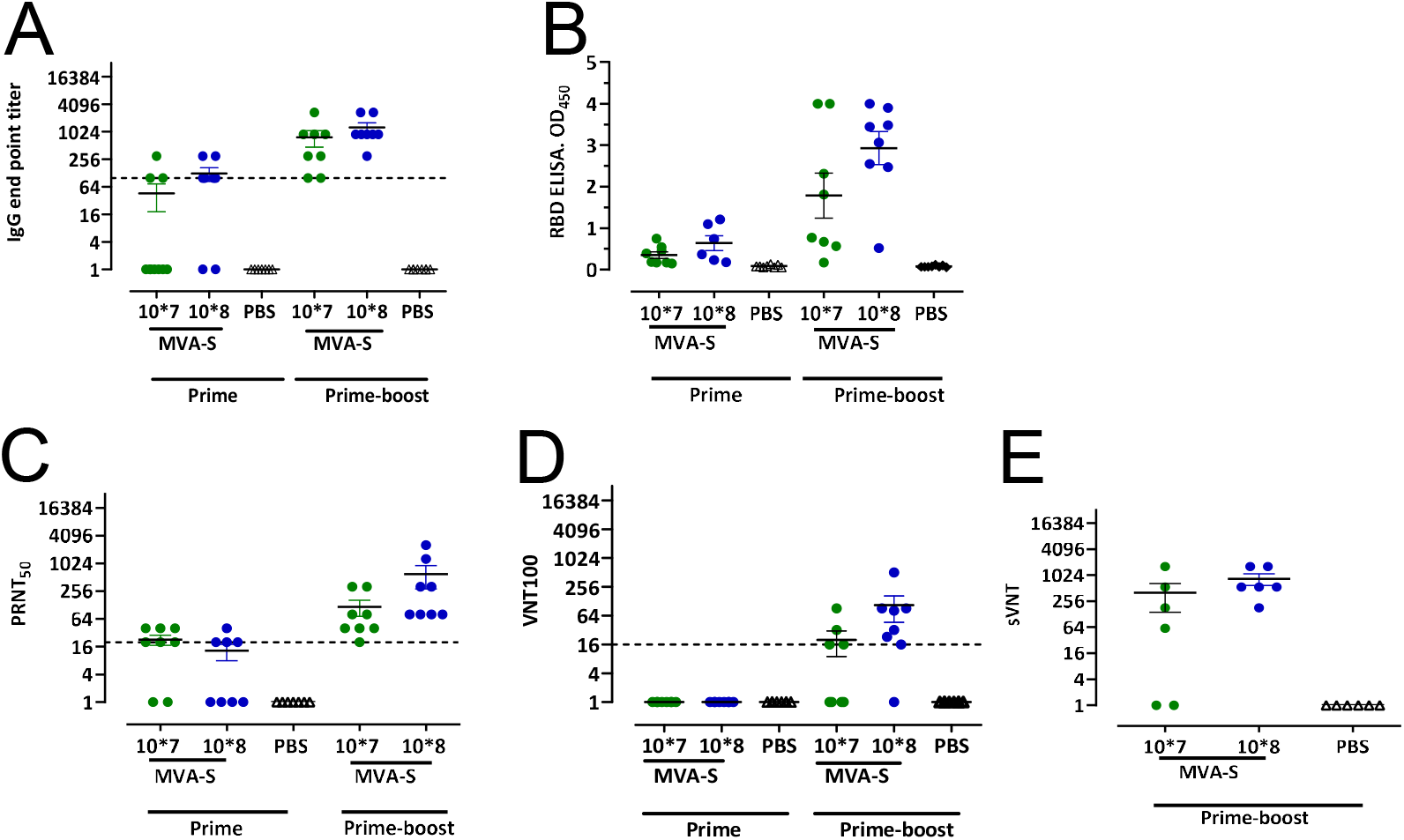
Antigen-specific humoral immunity induced by inoculation with recombinant MVA-SARS-2-S (MVA-S). Groups of BALB/c mice (n =7 to 12) were vaccinated in a prime-boost regime (21-day interval) with 10*7 or 10*8 PFU of MVA-S via the intra muscular (i.m) route. Mice inoculated with saline (PBS) served as controls. **(A,B)** Sera were collected 18 days after the first immunization (prime) and 14 days after the second immunization (primeboost) and analyzed for SARS-2-S specific IgG titers by ELISA, and **(C-E)** SARS-CoV-2 neutralizing antibodies by plaque reduction assay (PRNT_50_), virus neutralization (VNT_100_) or surrogate virus neutralization test (sVNT).

### MVA-SARS-2-S induced T cell responses in mice

To assess the activation of SARS-CoV-2-specific cellular immunity, we monitored S-specific CD8+ and CD4+ T cells in BALB/c mice vaccinated with LD or HD MVA-SARS-2-S in prime and prime-boost immunization schedules using 3-week intervals (*SI Appendix*, Fig. S2). To assess S antigen-specific T cells by IFN-γ ELISPOT, we isolated splenocytes at day 8 after MVA-SARS-2-S prime or boost immunization and used S-specific peptide stimulation for activation upon *in vitro* culture. Since information is limited on antigen specificities of SARS-CoV-2-specific T cells, we screened the Immune Epitope Database (IEDB) to select putative S-specific peptide epitopes compatible with activation of CD8+ or CD4+ T cells (*SI Appendix*, Tables S1 and S2). When testing pools of the predicted peptides with splenocytes from BALB/c mice immunized with 10^8^ PFU MVA-SARS-2-S, we detected responses above background in several peptide pools and identified the immunodominant SARS-CoV-2 S H2-K^d^ epitope S_269-278_ (GYLQPRTFL; S1 N-terminal domain, *SI Appendix*, Fig. S4). To evaluate the primary activation of SARS-2-S epitope specific CD8+ T cells, we inoculated BALB/c mice once with LD or HD MVA-SARS-2 and analyzed splenocytes on day 8 after vaccination. Single intramuscular immunizations with MVA-SARS-2-S already induced substantial levels of S_269-278_ epitope-specific activated CD8+ T cells with mean numbers of 341 IFN-γ spot forming cells (SFC) in 10^6^ splenocytes for LD and 275 SFC for HD compared to control mice immunized with non-recombinant MVA (no SFC detectable) (Fig. 4*A*). ELISPOT data aligned well with FACS analysis of T cells intracellulary stained for IFN-γ, where we also found higher frequencies (means of 0.32-0.36%) and higher absolute numbers of IFN-γ+ CD8+ T cells in splenocytes from vaccinated animals compared to control mice (Fig. 4*B*). Substantial numbers of the activated IFN-γ+ CD8+ T cells also co-expressed TNF-α (means of 0.22% and 0.27% of TNF-α expressing cells from total IFN-γ+ expressing cells) (Fig. 4*C*). Of note, the magnitude of SARS-2-S-specific CD8+ T cells did not significantly differ when comparing the groups of mice immunized with LD or HD of MVA-SARS-2-S.

**Fig. 4.**
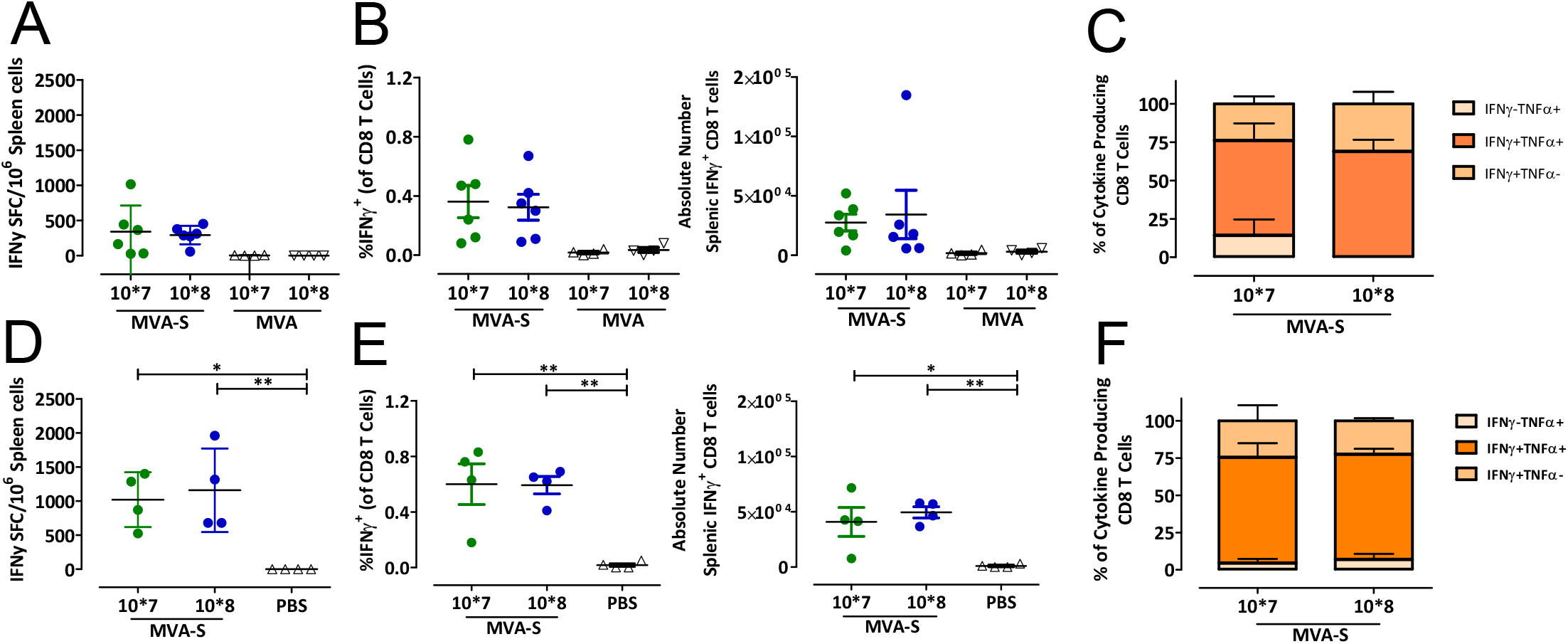
Activation of SARS-2-S specific CD8+ T cells after prime-boost immunization with MVA-SARS-2-S. Groups of BALB/c mice (n = 4-6) were immunized twice with 10*7 or 10*8 PFU of MVA-SARS-2-S (MVA-S) over a 21-day interval via the i.m. route. Mock immunized mice (PBS) were negative controls. **(A, B, C)** Splenocytes were collected and prepared on day 8 after prime, **(D, E, F)** or prime-boost immunization. Total splenocytes were stimulated with the H2d restricted peptide of the SARS-2-S protein S_268-276_ (S1; GYLQPRTFL) and were measured by IFN-γ ELISPOT assay and IFN-γ and TNF-α ICS plus FACS analysis. **(A, D)** IFN-γ SFC for stimulated splenocytes measured by ELISPOT assay. **(B, E)** IFN-γ production by CD8 T cells measured by FACS analysis. Graphs show the frequency and absolute number of IFN-γ+ CD8+ T cells. **(C, F)** Cytokine profile of S1-specific CD8 T cells. Graphs show the mean frequency of IFN-γ-TNF-α+, IFN-γ+TNF-α+ and IFN-γ+TNF-α-cells within the cytokine positive CD8+ T cell compartment. Differences between groups were analyzed by one-way ANOVA and Tukey post-hoc test. Asterisks represent statistically significant differences between two groups. * p < 0.05, ** p < 0.01

The booster immunizations on day 21 further increased the magnitudes of S-specific CD8+ T cells in response to MVA-SARS-2-S vaccination. At day 8 post boost, ELISPOT analysis revealed means of 1,020 IFN-γ SFC in LD vaccinated mice and 1,159 IFN-γ SFC in animals receiving HD MVA-SARS-2-S (Fig. 4*D*). Intracellular FACS analysis identified frequencies of 0.62% or 0.60% and absolute numbers of 40,873 or 49,553 IFN-γ+ CD8+ T cells for mice immunized with LD or HD MVA-SARS-2-S (Fig. 4*E*). Again, we confirmed that the majority (~75%) of IFN-γ+ CD8+ T cells also expressed TNF-α (Fig. 4*F*). The MVA-specific immunodominant CD8+ T cell determinant F2(G)_26-34_ (SPGAAGYDL(26)) served as a control peptide for the detection and comparative analysis of MVA vector-specific CD8+ T cells in BALB/c mice (*SI Appendix*, Fig. S5 and Fig. S6). In addition, using SARS-2-S derived peptides with predicted capacity for MHC II binding we also monitored the presence of activated CD4+ T cells. Using three different peptide pools (*SI Appendix*, Table S2), we confirmed the presence of spike-specific CD4+ T cells in the spleens of mice immunized with LD and HD prime-boost regiments (*SI Appendix*, Fig. S7).

### Protective capacity of MVA-SARS-2-S upon SARS-CoV-2 challenge

To model productive infection with SARS-CoV-2, we used an adenoviral transduction-based mouse model similar to those described recently (27, 28). We intratracheally transduced MVA-SARS-2-S vaccinated BALB/c mice with 5×10^8^ PFU of an adenoviral vector encoding both the human ACE2 receptor and the marker protein mCherry (ViraQuest Inc., North Liberty, IA, USA) at about two weeks after prime-boost immunization. Three days later the animals were infected with 1.5×10^4^ tissue culture infectious dose 50 (TCID_50_) SARS-CoV-2 (isolate BavPat1/2020 isolate, European Virus Archive Global # 026V-03883), and four days post challenge the animals were sacrificed, blood samples taken, and the lungs harvested to measure viral loads. Substantial virus RNA loads, on average >1,000 SARS-CoV-2 genome equivalents/ng of total RNA, were found in mock-immunized control mice. In contrast, the lung tissue of both LD and HD MVA-SARS-2-S-immunized animals contained significantly lower levels of SARS-CoV-2 RNA (<100 genome equivalents/ng total RNA; Fig. 5*A*). Adenoviral vector transduction levels of lung tissues were analyzed by real-time RT-PCR analysis to confirm comparable amounts of mCherry RNA (Fig. 5*B*). In addition, we found easily detectable levels of infectious SARS-CoV-2 (>1000 TCID_50_/ml) in the lungs from control but not in the tissues of immunized mice indicating the efficient inhibition of SARS-CoV-2 replication by vaccine-induced immune responses (Fig. 5*C*). In agreement with these data, only sera from MVA-SARS-2-S vaccinated animals (10/11) contained SARS-CoV-2 neutralizing circulating antibodies (Fig. 5*D*).

**Fig. 5.**
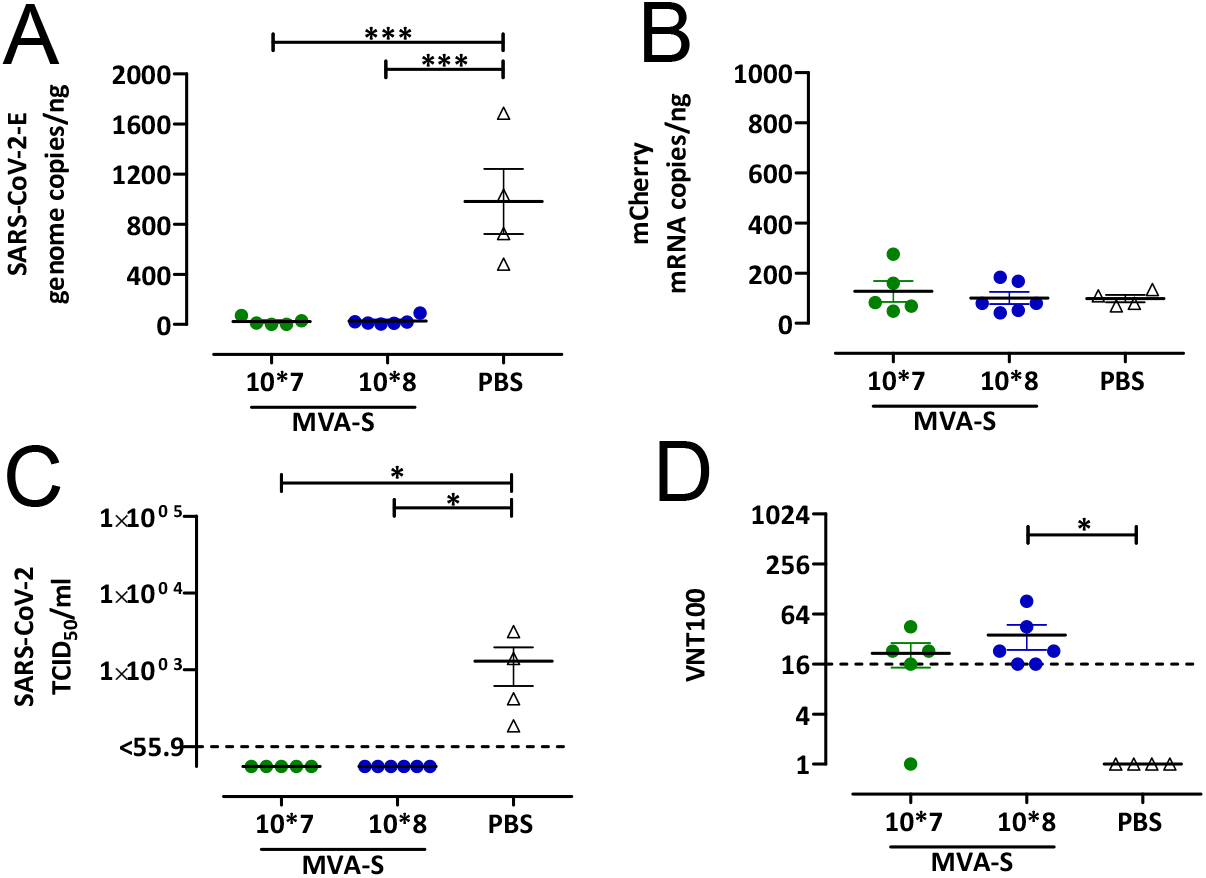
Protective capacity of MVA-SARS-2-S immunization against SARS-CoV-2 infection in human ACE2-transduced BALB/c mice. Groups of BALB/c mice (n = 4-6) were immunized twice with 10*7 or 10*8 PFU of MVA-SARS-2-S (MVA-S) over a 21-day interval via the i.m. route. Mock immunized mice (PBS) served as controls. About two weeks after the last immunization the mice were sensitized with an adenovirus expressing hACE2 and mCherry and infected with SARS-CoV-2 five days after transduction. Four days post challenge the animals were sacrificed and samples were taken for further analysis. **(A)** Lung tissues were harvested to determine SARS-CoV-2 RNA loads by viral genome copies, (B) the expression of mCherry by mRNA copies, or **(C)** the amounts of infectious SARS-CoV-2 by TCID_50_/ml. **(D)** Sera were tested for SARS-CoV-2 neutralizing antibodies by virus neutralization (VNT_100_). Statistical evaluation was performed with GraphPad Prism for Windows. Statistical significance of differences between groups is indicated as follows: *, *p* < 0.05; ***, *p* < 0.001.

## Discussion

Here, we report that the COVID-19 candidate vaccine MVA-SARS-2-S is compatible with clinical use and industrial-scale production. Building on extensive prior experience developing a candidate vaccine against MERS (17–19, 21), we selected the full-length SARS-CoV-2 S protein for delivery by recombinant MVA. The vector virus replicated efficiently in DF-1 cells, the cell substrate for an optimized manufacturing process, and MVA-SARS-2-S stably produces S glycoprotein antigen upon serial amplifications at low multiplicities of infection.

Similar to our experience gained with MVA-MERS-S, expression of the SARS-CoV-2 S gene by recombinant MVA resulted in a glycoprotein migrating at a molecular mass of about 190 kDa. Treatment with glycosidase to remove all N-linked carbohydrates produced a polypeptide of 145 kDa, closely corresponding to the molecular weight predicted from S gene nucleotide sequence. In addition, we observed S1 and S2 cleavage of the full-length SARS-2 S polypeptide, apparently with various efficiencies of proteolytic processing depending on the cell substrate of protein production. This finding is in agreement with previous reports suggesting complex activation of the betacoronavirus S glycoproteins, including the involvement of multiple cleavage events and several host proteases (29, 30). Similar to our findings with the MVA-produced MERS-CoV S protein, SARS-CoV-2 S-specific detection by immunofluorescence included strong surface staining of MVA-SARS-2-S infected cells. We conclude that the recombinant SARS-CoV-2 S protein is trafficking through the Golgi apparatus and is expressed at the cell surface, as shown previously for functional S produced from plasmid expression vectors (7, 31, 32).

Since the biochemical characterization of the MVA-expressed S suggested production of a mature and properly folded spike antigen, we investigated whether MVA-SARS-2-S elicits S-specific immune responses. In proof-of-principle experiments, mice receiving the MVA-SARS-2-S vaccine twice intramuscularly developed circulating S-specific antibodies that neutralized SARS-CoV-2 infections in cell culture and mounted high levels of SARS-2-S-specific CD8+ T cells. MVA-SARS-2-S elicited levels of virus neutralizing antibodies in BALB/c mice that were comparable to those induced by ChAdOx1nCoV-19 or MVA-MERS-S vaccinations (18, 33) and evidence from preclinical studies in non-human primates and hamsters indicate that vaccine-induced SARS-CoV-2 neutralizing antibodies correlate with protection against lung infection and clinical disease (34–36). The humoral immune responses elicited by MVA-SARS-2-S and measured by ELISA, two different SARS-CoV-2 neutralization assays and a surrogate neutralization assay pointed to a clear benefit of a booster immunization. These data are in line with results from phase 1 clinical testing of our MVA-MERS-S candidate vaccine providing evidence of humoral immunogenicity using homologous prime-boost vaccination (21). For SARS-CoV-2 neutralizing antibodies we found a strong correlation between the results obtained from authentic virus neutralization (PRNT_50_, VNT_100_) and the data from surrogate neutralization (sVNT). These data corroborate the findings of a recent study comparing this high-throughput sVNT assay to a pseudotyped virus neutralization assay based on SARS-CoV-2 S protein-carrying vesicular stomatitis virus (25).

The i.m. immunizations of BALB/c mice with low and high-dose MVA-SARS-2-S induced robust and nearly equal amounts of SARS-S-specific CD8+ T cells in prime and prime-boost vaccination. The average number of S-specific T cells was comparable to the average number of MVA vector-specific T cells highlighting the strong immunogenicity of MVA-SARS-2-S for inducing a SARS-S-specific CD8+ T cell response. The importance of vaccine induced T cell responses is illustrated by studies not only monitoring the adaptive immunity to SARS-CoV-2 in patients, but also demonstrating that strong SARS-CoV-2-specific CD4+ or CD8+ T cell responses are associated with low disease severity in individuals with COVID-19 (5).

When tested in a mouse model of SARS-CoV-2 lung infection all vaccinated BALB/c mice exhibited little or no replication of SARS-CoV-2, irrespective whether low or high-dose MVA-SARS-2-S was used for vaccination. Particularly encouraging was the complete absence of detectable infectious virus in the lung tissues of immunized animals. Notably, we found no evidence of a potential enhancement of SARS-CoV-2 infection through S-antigen-specific antibody induction, confirming our data with MVA-MERS-S that the S glycoprotein is an important and safe vaccine antigen (18, 19).

Overall, the MVA-SARS-2-S vector vaccine merits further development and the results presented here provided information for the start of a phase 1 clinical trial on 30 September 2020. To counteract the SARS-CoV-2 pandemic, candidate vaccines are being rapidly investigated in unprecedented numbers, and first front-runner vaccines are obtaining emergency licensing in Europe and the USA in 2020 (37). Yet, there is still much to learn when moving forward in COVID-19 vaccination. We expect that optimized protective immunity to COVD-19 will require vaccine approaches eliciting antiviral SARS-CoV-2-specific CD4+ and CD8+ T cells in a coordinated manner, together with virus neutralizing antibodies, in various population groups including children, the elderly and individuals with comorbidities.

## Materials and Methods

### Cell cultures

DF-1 cells (ATCC^®^ CRL-12203™) were maintained in VP-SFM medium (Thermo Fisher Scientific, Planegg, Germany), 2% heat-inactivated fetal bovine serum (FBS) (Thermo Fisher Scientific, Planegg, Germany) and 2% L-glutamine (Thermo Fisher Scientific, Planegg, Germany). Primary chicken embryonic fibroblasts (CEF) were prepared from 10 to 11-day-old chicken embryos (SPF eggs, VALO, Cuxhaven, Germany) using recombinant trypsin (Tryple TM, Thermo Fisher Scientific, Planegg, Germany) and maintained in VP-SFM medium, 10% FBS and 1% L-glutamine. Vero cells (ATCC CCL-81) were maintained in Dulbecco`s Modified Eagle’s Medium (DMEM), 10% FBS and 1% MEM non-essential amino acid solution (Sigma-Aldrich, Taufkirchen, Germany). Human A549 cells (ATCC^®^ CCL-185™) (LGC standards) were maintained in DMEM with high glucose and 10% FBS. Human HeLa cells (ATCC CCL-2) were maintained in Minimum Essential Medium Eagle (MEM) (Sigma-Aldrich, Taufkirchen, Germany), 7% FBS and 1% MEM non-essential amino acid solution. Human HaCat cells (CLS Cell Lines Service GmbH, Eppelheim, Germany) were maintained in DMEM, 10% FBS, 1% MEM non-essential amino acid solution and 1% HEPES solution (Sigma-Aldrich, Taufkirchen, Germany). All cells were cultivated at 37 °C and 5 % CO_2_.

### Plasmid construction

The coding sequence of the full-length SARS-CoV-2 S protein (SARS-2-S) was modified *in silico* by introducing silent mutations to remove runs of guanines or cytosines and termination signals of vaccinia virus-specific early transcription. In addition, a C-terminal tag sequence encoding nine amino acids (YPYDVPDYA, aa 98-106(38) from influenza virus hemagglutinin (HAtag) was added. The modified SARS-2-S cDNA was produced by DNA synthesis (Eurofins, Ebersberg, Germany) and cloned into the MVA transfer plasmid pIIIH5red under transcriptional control of the synthetic vaccinia virus early/late promoter PmH5 (22) to obtain the MVA expression plasmid pIIIH5red-SARS-2-S.

### Generation of recombinant viruses

MVA vector viruses were obtained following the established protocols for vaccine development as described in previous studies (17, 21, 39) MVA (clonal isolate MVA-F6-sfMR) was grown on CEF under serum-free conditions and served a as non-recombinant backbone virus to construct MVA vector viruses expressing the SARS-CoV-2 S gene sequences. Briefly, monolayers of 90-95% confluent DF-1 or CEF cells were grown in six-well tissue culture plates (Sarstedt, Nürnbrecht, Germany), infected with non-recombinant MVA at 0.05 multiplicity of infection (MOI), and transfected with plasmid pIIIH5red-SARS-2-S DNA using X-tremeGENE HP DNA Transfection Reagent (Roche Diagnostics, Penzberg, Germany) according to the manual. Afterwards, cell cultures were collected and recombinant MVA viruses were clonally isolated by serial rounds of plaque purification on DF-1 or CEF cell monolayers monitoring for transient co-production of the red fluorescent marker protein mCherry. To obtain vaccine preparations, recombinant MVA–SARS-2-S were amplified on CEF or DF-1 cell monolayers grown in T175 tissue culture flasks, purified by ultracentrifugation through 36% sucrose and reconstituted to high titre stock preparations in Tris-buffered saline pH 7.4. Plaque-forming units (PFU) were counted to determine viral titres (23).

### In vitro characterization of recombinant MVA-SARS-2-S

Genetic identity and genetic stability of vector viruses was confirmed by polymerase chain reaction (PCR) using viral DNA and detection of S-protein synthesis following serial passage at low MOI. For the latter, 95% confluent DF-1 cells were infected at MOI 0.05, incubated for 48h, harvested and used for reinfection. In total, five rounds of low MOI passage were performed. After the fifth passage, sixty virus isolates were obtained and amplified in 24-well DF-1 cultures for further testing. PCR analysis was performed to confirm genetic stability of viral genomes and MVA- and SARS-2-S-specific immunostaining served to monitor recombinant gene expression. The replicative capacity of recombinant MVA was tested in multi-step-growth experiments on monolayers of DF-1, HaCat, HeLa or A549 cells grown in 6-well-tissue-culture plates. Viruses were inoculated at MOI 0.05, harvested at 0, 4, 8, 24, 48, and 72 h after infection, and titrated on CEF monolayers to determine infectivities in cell lysates in PFU.

### Western Blot analysis of recombinant protein

To monitor production of the recombinant SARS-2-S protein, DF-1 cells were infected at MOI 10 with recombinant or non-recombinant MVA or remained uninfected (mock). At indicated time points of infection, cell lysates were prepared from infected cells and stored at −80 °C. Proteins from lysates were separated by electrophoresis in a sodium dodecyl sulfate (SDS)-10% polyacrylamide gel (SDS-PAGE; Bio-Rad, Munich) and subsequently transferred to a nitrocellulose membrane by electroblotting. The blots were blocked in a phosphate buffered saline (PBS) buffer containing 5% Bovine Serum Albumin (BSA) (Sigma-Aldrich, Taufkirchen, Germany) and 0.1% Tween-20 (Sigma-Aldrich, Taufkirchen, Germany) and incubated for 60 min with primary antibody, monoclonal anti-HAtag antibody (1:8000; HA Tag mAb 2-2.2.14, Thermo Fisher Scientific, Planegg, Germany) or COVID-19 patient serum (1:200). Next, membranes were washed with 0.1% Tween-20 in PBS and incubated with anti-mouse or anti-human IgG (1:5000; Agilent Dako, Glostrup, Denmark), conjugated to horseradish peroxidase. Blots were washed and developed using SuperSignal^®^ West Dura Extended Duration substrate (Thermo Fisher Scientific, Planegg, Germany). Chemiluminescence was visualized using the ChemiDoc MP Imaging System (Bio-Rad, Munich, Germany). For use of patient serum ethical approval was granted by the Ethics Committee at the Medical Faculty of LMU Munich (vote 20-225 KB) in accordance with the guidelines of the Declaration of Helsinki.

### Immunostaining of recombinant SARS-2-S protein

Vero cells were infected with 0.05 MOI MVA-SARS-2-S or non-recombinant MVA and incubated at 37 °C. After 24 h, cells were fixed with 4% paraformaldehyde (PFA) for 10 min on ice, washed two times with PBS, and permeabilized with 0.1% Triton X-100 (Sigma-Aldrich, Taufkirchen, Germany) solution in PBS. Permeabilized cells were probed with a monoclonal antibody against the HA-tag epitope (1:1000; HA Tag mAb 2-2.2.14, Thermo Fisher Scientific, Planegg, Germany) to detect SARS-2-S protein. Non-permeabilized cells were stained with a mouse monoclonal antibody obtained against the S protein of SARS-CoV-1 (SARS-1-S; 1:200; GeneTex) before fixation with PFA. Polyclonal goat anti-mouse secondary antibody (1:1000; Life Technologies, Darmstadt, Germany) was used to visualize S-specific staining by red fluorescence. Nuclei were stained with 1 μg/ml of 4,6-diamidino-2-phenylindole (DAPI) (Sigma-Aldrich, Taufkirchen, Germany) and cells were analyzed using the Keyence BZ-X700 microscope (Keyence, Neu-Isenburg, Germany) with a ×100 objective.

### Vaccination experiments in mice

Female BALB/c mice (6 to 10 week-old) were purchased from Charles River Laboratories (Sulzfeld, Germany). Mice were maintained under specified pathogen-free conditions, had free access to food and water, and were allowed to adapt to the facilities for at least one week before vaccination experiments were performed. All animal experiments were handled in compliance with the European and national regulations for animal experimentation (European Directive 2010/63/EU; Animal Welfare Acts in Germany). Immunizations were performed using intramuscular applications with vaccine suspension containing either 10^7^ or 10^8^ PFU recombinant MVA-SARS2-S, non-recombinant MVA or PBS (mock) into the quadriceps muscle of the left hind leg. Blood was collected on days 0, 18, 35, 56 or 70. Coagulated blood was centrifuged at 1300×g for 5 min in MiniCollect vials (Greiner Bio-One, Alphen aan den Rijn, The Netherlands) to separate serum, which was stored at −20 °C until further analysis.

### Transduction of vaccinated mice with Ad_ACE2-mCherry and challenge infection with SARS-CoV-2

All animal experiments were performed in accordance with Animal Welfare Acts in Germany and were approved by the regional authorities. Vaccinated mice were housed under pathogen-free conditions and underwent intratracheal inoculation with 5×10^8^ PFU Adenovirus-ACE2-mCherry (cloned at ViraQuest Inc., North Liberty, IA, USA) under ketamine/xylazine anesthesia. Three days post transduction, mice were infected via the intranasal route with 1.5×10^4^ tissue culture infectious dose 50 (TCID_50_) SARS-CoV-2 (BavPat1/2020 isolate, European Virus Archive Global # 026V-03883). Mice were sacrificed 4 days post infection and serum as well as lung tissue samples were taken for analysis of virus loads.

### Quantitative real-time reverse transcription PCR to determine SARS-CoV-2 or mCherry RNA

Tissue samples of immunized and challenged mice were excised from the left lung lobes and homogenized in 1 ml DMEM. SARS-CoV-2 titres in supernatants (in TCID_50_ per ml) were determined on VeroE6 cells. RNA isolation was performed with the RNeasy minikit (Qiagen) according to the manufacturer’s instructions. The RNA amount was measured using the NanoDrop ND-100 spectrophotometer. Total RNA was reverse transcribed and quantified by real-time PCR using the OneStep RT-PCR kit (Qiagen) as described previously(40) with the primer pair upE-Fwd and upE-Rev and the probe upE-Prb on a StepOne high-throughput fast real-time PCR system (ThermoFisher). Additionally, for every tissue sample from transduced and infected mice, evidence for successful ACE2 transduction was determined by real-time RT-PCR for mCherry mRNA with the OneStep RT-PCR kit (Qiagen). All samples for mCherry analysis were evaluated in one RT-PCR run. Quantification was carried out using a standard curve based on 10-fold serial dilutions of appropriate control RNA ranging from 10^2^ to 10^5^ copies.

### Antigen-specific IgG ELISA

SARS-2-S-specific serum IgG titres were measured by enzyme-linked immunosorbent assay (ELISA) as described previously(41). Flat bottom 96-well ELISA plates (Nunc MaxiSorp Plates, Thermo Fisher Scientific, Planegg, Germany) were coated with 50 ng/well recombinant 2019-nCoV (COVID-19) S protein (Full Length-R683A-R685A-HisTag, ACROBiosystems, Newark, USA) overnight at 4 °C. Plates were washed and then blocked for 1 h at 37 °C with blocking buffer containing 1% BSA (Sigma-Aldrich, Taufkirchen, Germany) and 0.15M sucrose (Sigma-Aldrich, Taufkirchen, Germany) dissolved in PBS. Mouse sera were serially diluted three-fold down the plate in PBS containing 1% BSA (PBS/BSA), starting at a dilution of 1:100. Plates were then incubated for 1 h at 37 °C. After incubating and washing, plates were probed with 100 μl/well of goat anti-mouse IgG HRP (1:2000; Agilent Dako, Denmark) diluted in PBS/BSA for 1 h at 37 °C. After washing, 100 μl/well of 3′3′, 5′5′-Tetramethylbenzidine (TMB) Liquid Substrate System for ELISA (Sigma-Aldrich, Taufkirchen, Germany) was added until a colour change was observed. The reaction was stopped by adding 100μl/well of Stop Reagent for TMB Substrate (450 nm, Sigma-Aldrich, Taufkirchen, Germany). Absorbance was measured at 450 nm. The absorbance of each serum sample was measured at 450 nm with a 620 nm reference wavelength. ELISA data were normalized using the positive control. The cut-off value for positive mouse serum samples was determined by calculating the mean of the normalized OD 450nm values of the PBS control group sera plus 6 standard deviations (mean + 6 SD).

### Surrogate virus neutralization assay (sVNT)

To test for the presence of neutralizing anti-SARS-CoV-2-S serum antibodies we used surrogate virus neutralization test as described before with slight modifications (25). Briefly, 6 ng of SARS-CoV-2 S RBD (Trenzyme) was pre-incubated for 1 hour at 37 °C with heat-inactivated test sera at final dilutions between 1:20 to 1:540, as indicated on the graphs. Afterwards, SARS-CoV-2 S RBD-serum mixtures were loaded onto MaxiSorp 96F plates (Nunc) coated with 200 ng/well ACE2 [produced in-house as described in Bosnjak et al.(25)] and blocked with 2% bovine serum albumin/2% mouse serum (Invitrogen) and incubated for additional 1 h at 37 °C. As controls we used SARS-CoV-2-S-RBD pre-incubated only with buffer and non-specific mouse serum (Invitrogen). Plates were extensively washed with phosphate-buffered saline/0.05% Tween-20 (PBST), followed by incubation for 1h at 37°C with an HRP-conjugated anti-His-tag antibody (1.2 μg/ml; clone HIS 3D5). After appropriate washing, colorimetric signals were developed by addition of the chromogenic substrate 3,3′,5,5′-tetramethylbenzidine (TMB; TMB Substrate Reagent Set, BD Biosciences) and stopped by addition of equal volume of 0.2 M H_2_SO_4_. The optical density values measured at 450nm and 570nm (SpectraMax iD3 microplate reader, Molecular Devices) were used to calculate percentage of inhibition after subtraction of background values as inhibition (%) = (1 - Sample OD value/Average SARS-CoV-2 S RBD OD value) x100. To remove background effects, the mean percentage of inhibition from non-specific mouse serum (Invitrogen) was deducted from sample values and neutralizing anti-SARS-CoV2-S antibodies titres were determined as serum dilution that still had binding reduction > mean + 2 SD of values from sera of vehicle-treated mice.

### Plaque reduction neutralization test 50 (PRNT_50_)

We tested serum samples for their neutralization capacity against SARS-CoV-2 (German isolate; GISAID ID EPI_ISL 406862; European Virus Archive Global #026V-03883) by using a previously described protocol(24). We 2-fold serially diluted heat-inactivated samples in Dulbecco modified Eagle medium supplemented with NaHCO3, HEPES buffer, penicillin, streptomycin, and 1% foetal bovine serum, starting at a dilution of 1:10 in 50 μL. We then added 50 μL of virus suspension (400 plaque-forming units) to each well and incubated at 37°C for 1 h before placing the mixtures on Vero-E6 cells. After incubation for 1 h, we washed, cells supplemented with medium, and incubated for 8 h. After incubation, we fixed the cells with 4% formaldehyde/phosphate-buffered saline (PBS) and stained the cells with polyclonal rabbit anti-SARS-CoV antibody (Sino Biological, https://www.sinobiological.com) and a secondary peroxidase-labeled goat anti-rabbit IgG (Dako, https://www.agilent.com). We developed the signal using a precipitate forming 3,3′,5,5′-tetramethylbenzidine substrate (True Blue; Kirkegaard and Perry Laboratories, https://www.seracare.com) and counted the number of infected cells per well by using an ImmunoSpot Image Analyzer (CTL Europe GmbH, https://www.immunospot.eu). The serum neutralization titre is the reciprocal of the highest dilution resulting in an infection reduction of >50% (PRNT_50_). We considered a titre >20 to be positive.

### SARS-CoV-2 virus neutralization test (VNT_100_)

The neutralizing activity of mouse serum antibodies was investigated based on a previously published protocol (9). Briefly, samples were serially diluted in 96-well plates starting from a 1:16 serum dilution. Samples were incubated for 1 h at 37°C together with 100 50% tissue culture infectious doses (TCID_50_) of SARS-CoV-2 (BavPat1/2020 isolate, European Virus Archive Global # 026V-03883). Cytopathic effects (CPE) on VeroE6 cells (ATCC CRL1586) were analyzed 4 days after infection. Neutralization was defined as the absence of CPE compared to virus controls. For each test, a positive control (neutralizing COVID-19 patient plasma) was used in duplicates as an inter-assay neutralization standard. Ethical approval was granted by the Ethics Committee at the Medical Faculty of LMU Munich (vote 20-225 KB) in accordance with the guidelines of the Declaration of Helsinki.

### Prediction and generation of synthetic SARS-2-S peptides

The sequence of the SARS-CoV-2 S protein (NCBI ID: QHD43416.1, Uniprot ID: P0DTC2 (SPIKE_SARS2)) served for epitope prediction, and probable CD8+ and CD4+ T cell determinants were examined with the Immune Epitope Database and Analysis Resource (IEDB, https://www.iedb.org/). For identification of potential CD8+ T cell determinants, the MHC-I Binding Prediction and MHC-I Processing Prediction tools (42, 43) were used and projections for 9-11mer peptides spanning the entire SARS-2-S protein sequence were obtained. The inputs selected for the search included the Prediction Method ‘IEDB recommended 2.22’, the MHC source species ‘Mouse’ and the MHC class I alleles H2-K^d^, H2-D^d^ and H2-L^d^. The output was restricted to a percentile rank cut-off of 10.0. After lists of peptides were generated, all peptides with an IC50 score of 500nM or less were selected for inclusion in the top 5% list. All the peptides in this list were further analyzed using the MHC-I Processing Prediction tool ‘Proteasomal cleavage/TAP transport/MHC class I combined predictor’. All peptides with an IC50 score of 500nM or less and a high total score were chosen and subsequently included in the top peptides list. To confirm that these peptides were potential binders of MHC class I alleles H2-K^d^, H2-D^d^ and H2-L^d^, they were further screened for MHC I binding using the RankPep server(44). Peptides that were found to bind to any of the above alleles were selected for synthesis and testing.

For the identification of potential CD4+ T cell determinants, the MHC-II Binding Prediction tool (42, 43) served to obtain 15mer peptides spanning the entire SARS-2-S protein sequence. The inputs for the analysis included the Prediction Method ‘IEDB recommended 2.22’, the MHC source species ‘Mouse’ and the MHC class II alleles H2-IA^d^ and H2-IE^d^. Peptides with percentile rank of 10.0 or less and an IC50 score of 1000 nM or less were further tested for MHC class II binding using the RankPep server (44). Peptides bound to any of the above MHC class II alleles were selected for synthesis and testing. All peptides were obtained from Thermo Fisher Scientific (Planegg, Germany) as crude material (<50% purity) at a 1–4 mg scale, dissolved in PBS or DMSO to 2 mg/ml, aliquoted and stored at −20 °C.

### T cell analysis by Enzyme-Linked Immunospot (ELISPOT)

At days 8 and 14 post prime or prime-boost vaccination, mice were sacrificed and splenocytes were prepared. Briefly, spleens were passed through a 70 μm strainer (Falcon^®^, Sigma-Aldrich, Taufkirchen, Germany) and incubated with Red Blood Cell Lysis Buffer (Sigma-Aldrich, Taufkirchen, Germany). Cells were washed and resuspended in RPMI-10 (RPMI 1640 medium containing 10% FBS, 1% Penicillin-Streptomycin, 1% HEPES; Sigma-Aldrich, Taufkirchen, Germany). ELISPOT assay (Mabtech ELISpot kit for mouse IFN-γ, Biozol, Eching, Germany) was performed to measure IFN-γ-producing T cells following the manufacturer’s instructions. Briefly, 2×10^5^ splenocytes/100μl were seeded in 96-well plates and stimulated with individual peptides (2 μg/mL RPMI-10). Non-stimulated cells and cells stimulated with phorbol myristate acetate (PMA) / ionomycin (Sigma-Aldrich, Taufkirchen, Germany) or vaccinia virus peptide SPGAAGYD (F2(G)_26-34_; H-2L^d^;(26)) served as controls. After incubation at 37 °C for 48 h, plates were stained according to the manufacturer’s instructions. Spots were counted and analyzed by using an automated ELISPOT plate reader and software following the manufacturer’s instructions (A.EL.VIS Eli.Scan, A.EL.VIS ELISPOTAnalysis Software, Hannover, Germany).

### T cell analysis by Intracellular Cytokine Staining (ICS)

The detailed methods for intracellular cytokine staining (ICS) were described previously(41). Briefly, whole splenocytes were diluted in RPMI-10 and plated onto 96-well-U-bottom plates using 10^6^ cells/well. Cells were stimulated with 8 μg/ml S_269-278_ peptide or vaccinia virus peptide F226-34 for analysis of SARS-2-S- or MVA-specific CD8+ T cells. Splenocytes stimulated with PMA (10 ng/ml) plus ionomycin (500 ng/ml) served as positive controls and RPMI alone was used as a negative control. After 2 h at 37 °C, brefeldin A (Biolegend, San Diego, CA, USA) was added according to the manufacturer’s instructions and stimulated cells were further maintained for 4 h at 37 °C. After the stimulations, cells were washed with FACS buffer (MACSQuant Running Buffer, Miltenyi Biotec, Bergisch Gladbach, Germany, plus 2% FBS) and stained extracellularly with anti-mouse CD3 phycoerithrin (PE)-Cy7 (clone 17A2, 1:100, Biolegend), anti-mouse CD4 Brilliant Violet 421 (clone GK1.5, 1:600, Biolegend), anti-mouse CD8α Alexa Fluor 488 (clone 53-6.8, 1:300, Biolegend), and purified CD16/CD32 (Fc block; clone 93, 1:500, Biolegend) using 50 μl/well diluted in FACS Buffer for 30 min on ice. After staining and washing, cells were incubated with 100 μl/well of the fixable dead cell viability dye Zombie Aqua (1:800, Biolegend) diluted in PBS for 30 min on ice. Cells were then washed, fixed with 100 μl/well of Fixation Buffer (Biolegend) for 20 min at room temperature, washed again, resuspended in 200 μl/well of FACS buffer and stored overnight at 4 °C. Next, cells were permeabilized using Intracellular Staining Permeabilization Wash Buffer (Perm Wash buffer; Biolegend; dilution 1:10), and stained intracellularly in 100 μl/well of anti-mouse IFN-γ (clone XMG1.2, 1:200, Biolegend) plus anti-mouse TNF-α (clone MP6-XT22, 1:200, Biolegend) diluted in Perm Wash buffer for 30 min at room temperature. Thereafter, cells were washed with Perm Wash buffer and resuspended in FACS buffer. Prior to analysis, samples were filtered through a 50 μm nylon mesh (Sefar Pty Ltd., Huntingwood, NSW, Australia) into 5 ml round bottom FACS tubes (Sarstedt, Nümbrecht, Germany). For each antibody, single colour controls were prepared using OneComp eBeads™ Compensation Beads (eBioscience, Thermo Fisher Scientific) and cells for the viability dye Zombie Aqua. Data was acquired by the MACSQuant VYB Flow Analyser (Miltenyi Biotec) and analyzed using FlowJo (FlowJo LLC, BD Life Sciences, Ashland, OR, USA).

### Statistical analysis

Data were prepared using GraphPad Prism version 5 (GraphPad Software Inc., San Diego CA, USA) and expressed as mean ± standard error of the mean (SEM). Data were analyzed by unpaired, two-tailed t-tests to compare two groups and one-way ANOVA to compare three or more groups. P < 0.05 was used as the threshold for statistical significance.

## Acknowledgments

We thank Patrizia Bonert, Ursula Klostermeier, Johannes Döring and Axel Groß for expert help in animal studies. We thank Nico Becker, Astrid Herwig, Lennart Kämpfer for help with BSL3 sample preparation and testing. This work was supported by the German Center for Infection Research (DZIF: projects TTU 01.921 to GS and SB, TTU 01.712 to GS), the Federal Ministry of Education and Research (BMBF 01KX2026 to GS and SB, BMBF 01KI20702 to GS and SB, ZOOVAC 01KI1718, RAPID 01KI1723G to AV; “NaFoUniMedCovid19“ FKZ: 01KX2021, Project “B-FAST” to RF). Deutsche Forschungsgemeinschaft, (DFG, German Research Foundation) Excellence Strategy EXC 2155 “RESIST” (Project ID39087428 to RF), by DFG grant SFB900-B1 (Projektnummer 158989968 to RF) and by funds of the state of Lower Saxony (14-76103-184 CORONA-11/20 to RF).

