## SupplementalData for "Immunogenicity and efficacy of the COVID-19 candidate vector vaccine MVA SARS 2 S in preclinical vaccination"

### *SI Appendix*

#### Fig. S1

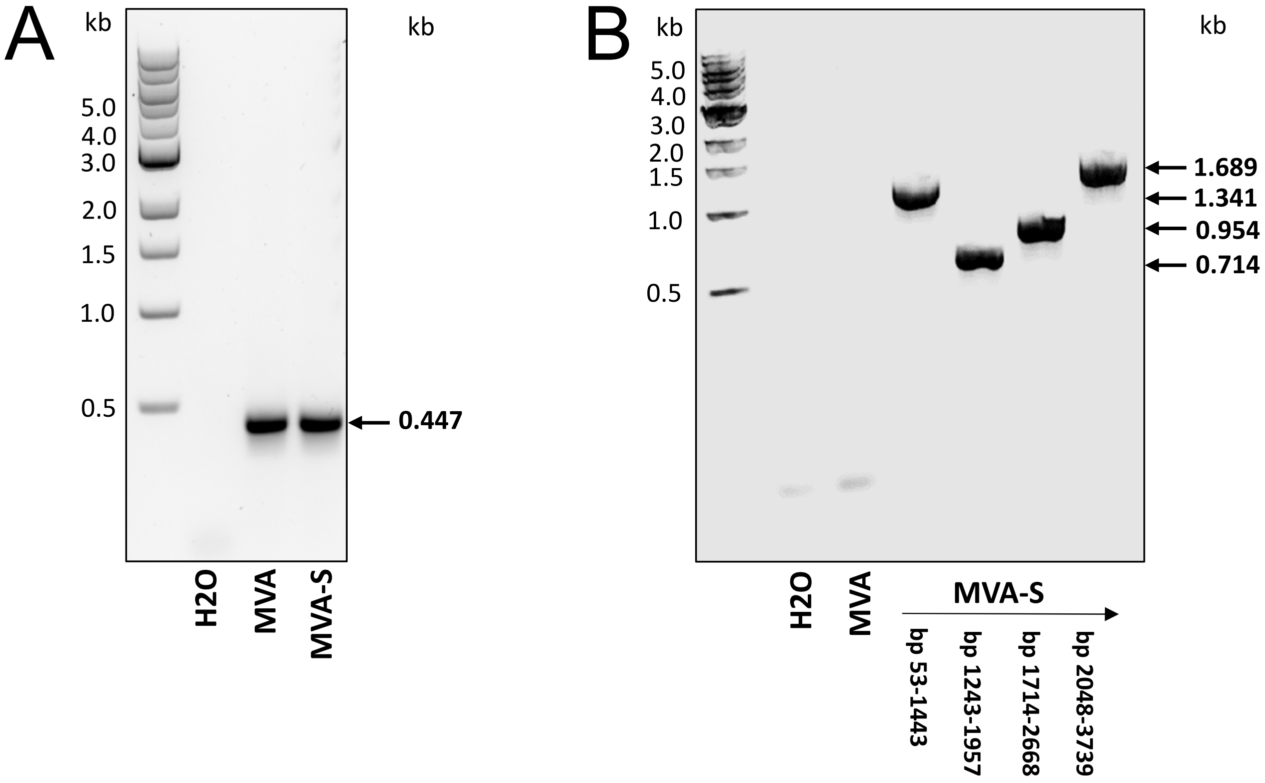


Suppl. Fig. S1. Molecular analysis of the MVA-SARS-2-S genome. PCR analysis of genomic viral DNA to monitor sequences of the *C7L* gene locus (**A**) and the SARS-CoV-2 S gene (**B**) in the MVA-SARS-2-S genome. **(A)** PCR amplification of a specific 447 bp DNA fragment from the MVA *C7L* gene sequence suggested the integrity of the *C7L* gene locus in the recombinant MVA-SARS-2-S genome (MVA-S) in comparison to the non-recombinant MVA genome (MVA). The *C7L* gene is non-essential for MVA growth in chicken fibroblast cultures but the gene function is necessary to maintain unimpaired expression of MVA or recombinant genes under transcriptional control of vaccinia virus-specific late promoters (Backes S et al., JGV 91:470-482. doi:10.1099/vir.0.015347-0). (**B**) Four different PCRs were used to assess the integrity of the full-length SARS-2-S gene sequence inserted in the MVA-SARS-2-S genome. Specifically amplified DNA fragments demonstrated the expected molecular weight with 1.341 kp (specific for S nucleotides 53-1443), 0.714 kb (specific for S nucleotides 1243-1957), 0.954 kb (specific for S nucleotides 1714-2668) and 1.689 kb (specific for S nucleotides 2048-3739) from the SARS-2 S gene sequence.

Fig. S1

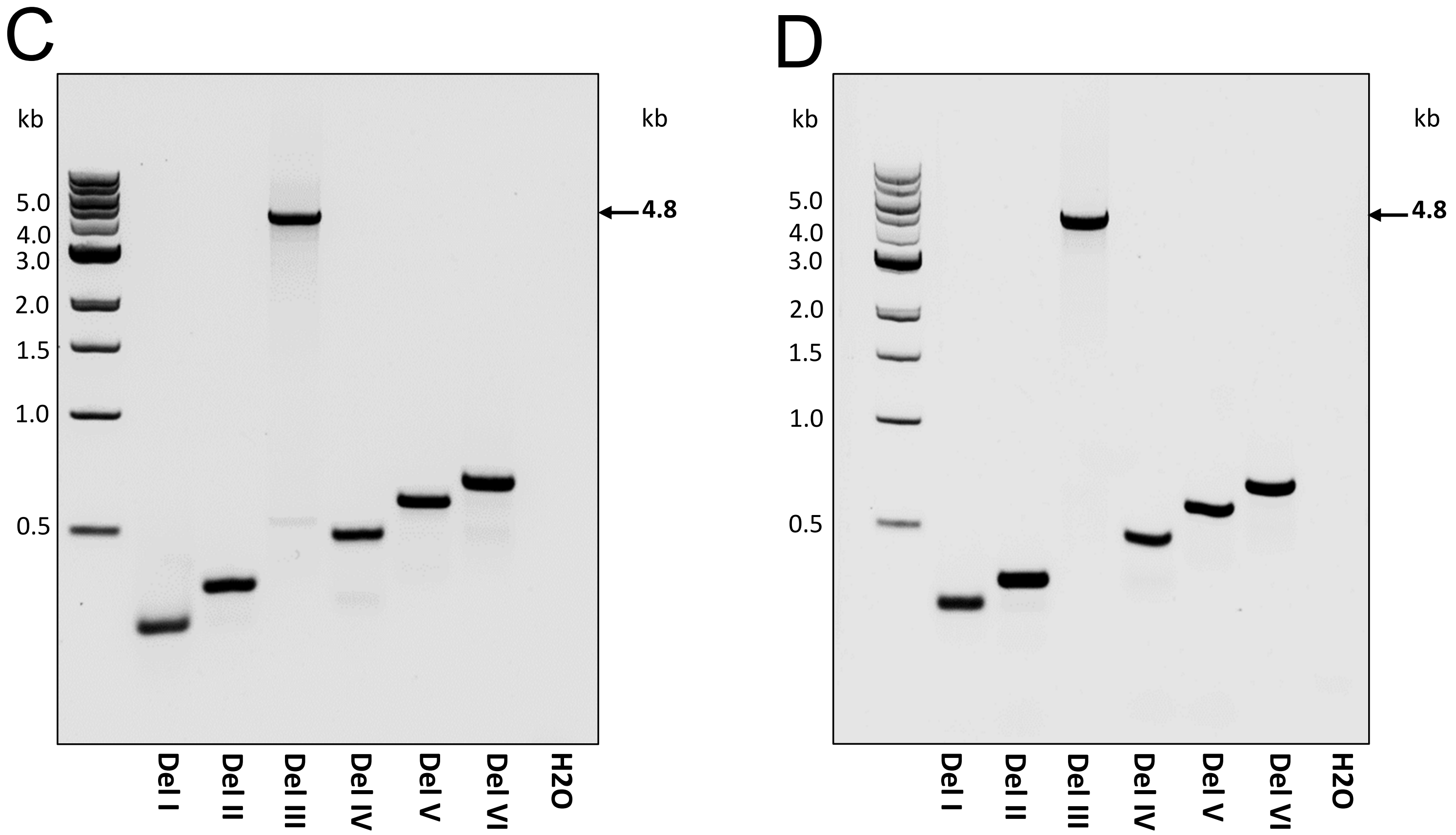


Suppl. Fig. S1. Genetic stability of MVA-SARS-2-S after serial DF-1 cell culture amplification. DF-1 cells were infected with MVA-SARS-2-S at a multiplicity of infection (MOI) of 0.05 and incubated for 48 h. Subsequently, the amplified virus was harvested and used to re-infect fresh DF-1 cells at MOI of 0.05 for 48 h. This procedure was performed five times. MVA-SARS-2-S genetic stability was tested by PCR analysis of genomic viral DNA and ~~the~~ monitoring for recombinant gene expression by S-specific immunostaining. PCR analysis demonstrated the genetic stability for six loci in the MVA-SARS-2-S genome (deletion sites Del I-VI) including the the heterologous SARS-CoV-2 S gene sequences inserted into the site of deletion III (Del III) with the amplification of characteristic size DNA fragments from viral DNA prepared after the first **(C),** or the fifth **(D)** round of MVA-SARS-2-S amplification in DF-1 cultures.

### Fig. S1

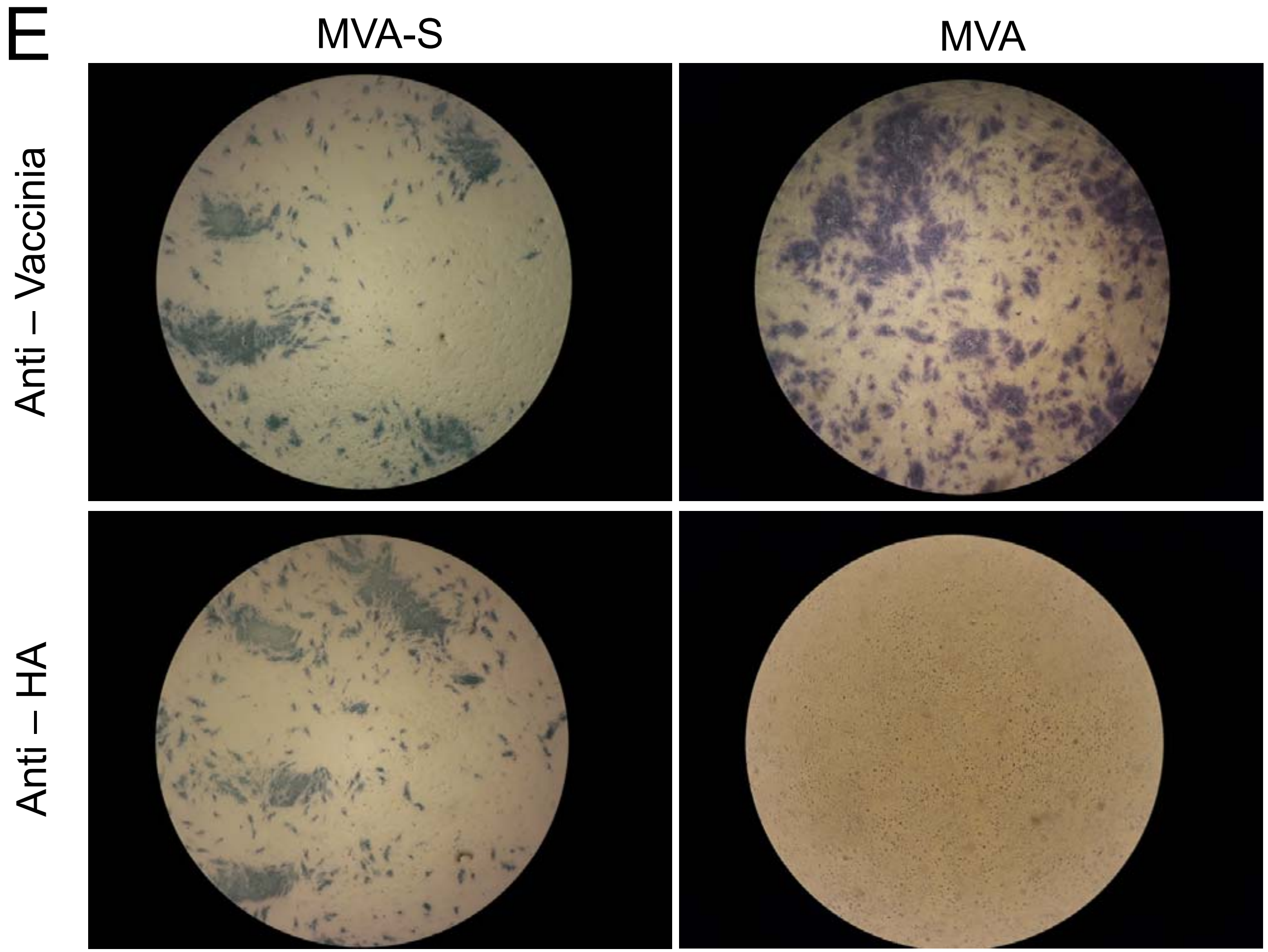


Suppl. Fig. S1. Unimpaired recombinant gene expression following serial DF-1 passage was tested by immunostaining for production of SARS-2-S using a mouse monoclonal antibody directed against the HA-tag (anti-HA). A total of 60 clonal MVA-SARS-2-S (MVA-S) isolates were picked after the fifth virus amplification in DF-1 cells and used to infect fresh DF-1 cell monolayers grown in 24-well tissue culture plates. Infections with non-recombinant MVA (MVA) served as controls. **(E)** After 48 h infection the cell monolayers were fixed and stained with anti-vaccinia and anti-HA antibody. All MVA-SARS-2-S isolates (60/60) tested positive for unimpaired expression of recombinant SARS-2-S.

Fig. S2

A

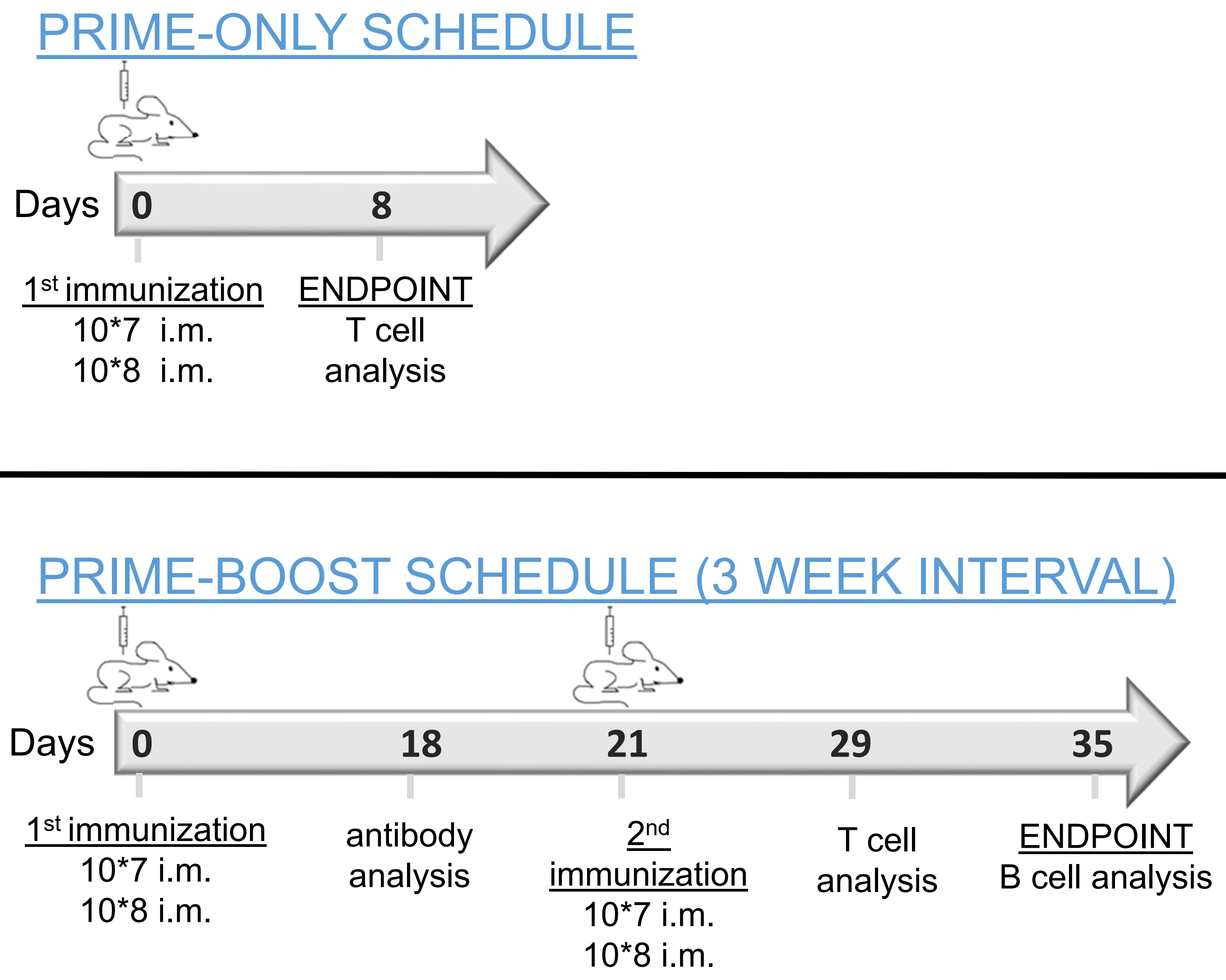
PRIME-ONLY SCHEDULE
Days
0
8
1st immunization
10*7 i.m.
10*8 i.m.
ENDPOINT
T cell
analysis
PRIME-BOOST SCHEDULE (3 WEEK INTERVAL)
Days
0
18
21
29
35
1st immunization
10*7 i.m.
10*8 i.m.
antibody
analysis
2nd
immunization
10*7 i.m.
10*8 i.m.
T cell
analysis
ENDPOINT
B cell analysis

### Fig. S2

B

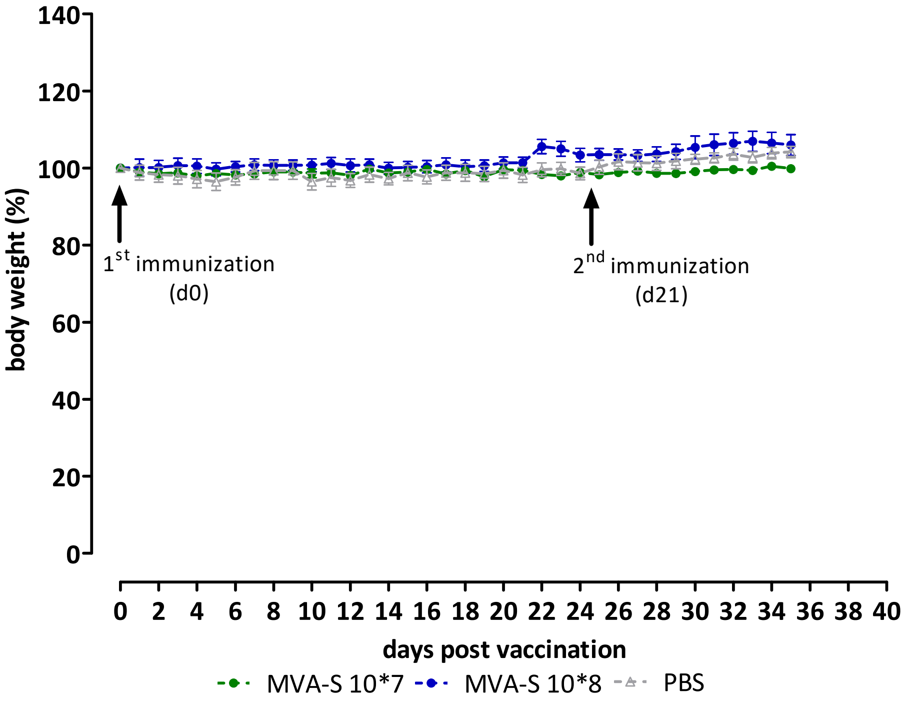


C

| Main groups | | | | | | | Sex: female |
|---|---|---|---|---|---|---|---|
| Group | | 1 | | | 2 | | |
| Treatment | | Vehicle | | | MVA-SARS-2-S prime boost | | |
| Dose | | | | | 10E8 pfu/animal | | |
| Day of necropsy | | 35 | | | 35 | | |
| Grading | | . | 1 | 2 | . | 1 | 2 |
| Organs: | Microscopic findings: | | | | | | |
| Adrenal gland | No finding(s) | 6/6 | 0/6 | 0/6 | 12/12 | 0/12 | 0/12 |
| Kidney, L | No finding(s) | 6/6 | 0/6 | 0/6 | 12/12 | 0/12 | 0/12 |
| Liver | Inflammatory cells, multifocal | 5/6 | 1/6 | 0/6 | 11/12 | 1/12 | 0/12 |
| Lungs | Prominent BALT | 6/6 | 0/6 | 0/6 | 11/12 | 1/12 | 0/12 |
| Ln. subiliacus, L | Lymphoid hyperplasia | 6/6 | 0/6 | 0/6 | 3/12 | 9/12 | 0/12 |
| Ln. ischiadicus, L | Lymphoid hyperplasia | 6/6 | 0/6 | 0/6 | 8/12 | 4/12 | 0/12 |
| Ln. popliteus, L | Lymphoid hyperplasia | 6/6 | 0/6 | 0/6 | 1/12 | 11/12 | 0/12 |
| Ln. iliacus medialis, L | Lymphoid hyperplasia | 6/6 | 0/6 | 0/6 | 1/12 | 11/12 | 0/12 |
| Thymus | No finding(s) | 6/6 | 0/6 | 0/6 | 12/12 | 0/12 | 0/12 |
| Spleen | No finding(s) | 6/6 | 0/6 | 0/6 | 12/12 | 0/12 | 0/12 |
| Site of administration, L | Mixed cell infiltration, interstitial, multifocal | 6/6 | 0/6 | 0/6 | 3/12 | 8/12 | 1/12 |
| Site of administration, L | Necrosis, myofibres | 6/6 | 0/6 | 0/6 | 10/12 | 2/12 | 0/12 |
| Site of administration, L | Degeneration, myofibres | 2/6 | 4/6 | 0/6 | 0/12 | 12/12 | 0/12 |

### No. of animals affected / total No. of animals.

Fig. S2. MVA-SARS-2-S immunization schedules and monitoring for side effects of vaccination. Groups of BALB/c mice (n=6-12) were vaccinated with low dose 10*7 PFU (LD) or high dose 10*8 PFU (HD) MVA-SARS-2-S via the intramuscular route using two different schedules (prime-only n=6, prime-boost 3 week interval, n=12). **(A)** Schematic diagram of the immunization schedules. T cell responses were tested at day 8 post 1st immunization (prime) or 2nd immunization (boost). B cell responses (including S-binding and SARS-CoV-2 neutralizing antibodies) were examined three days before the 2nd immunization (d18 vs. d53) and 14 days post 2nd immunization (d35). **(B)** Monitoring for body weight changes of mice after prime-boost vaccination with MVA-SARS-2-S. Vaccination with saline (PBS) was used as a control. Body weights were measured daily. No signs of discomfort or disease were observed in vaccinated or control animals. **(C)** Histopathological examinations in prime-boost vaccinated animals. L = Left side. No lesions could be attributed to MVA-SARS-S-2 inoculation in any tissue other than the administration site and draining lymph nodes. Systemic effects related to vaccination were not seen. Signs of minimal to mild myodegeneration at the administration site were observed in treated and control mice. Local inflammation of the myofiber interstitium and the adjacent adipose tissue was observed and interpreted as part of the physiological immune reaction to the vaccine virus as a consequence of the treatment procedure. The degree and extent of inflammation, myodegeneration and necrosis was in accordance with the ratio of inoculum volume in relation to the administration site. The lymphoid hyperplasia observed in draining lymph nodes is interpreted as a sign of immune competence of the animals and is characteristic for any early response to inflammation at a draining site. In conclusion, we observed no evidence for ~~a~~ potential toxicity of the full human dose of MVA-SARS-2-S in BALB/c mice. The repeated vaccination was well tolerated and caused no adverse events and no relevant macroscopic or histopathological changes. The observed reactions were comparable to previous experiments using non-recombinant MVA or other recombinant MVA vaccine constructs and are considered to be part of the pharmacodynamic principle of MVA-based vaccination. (Langenmayer et al. Distribution and absence of generalized lesions in mice following single dose intramuscular inoculation of the vaccine candidate MVA-MERS-S. Biologicals. 2018;54:58-62.)

### Table S1

Table S1. Selected SARS-CoV-2-S peptides with predicted MHC class I (H2d) restriction

| Peptide ID | Peptide | Length | Start | End | Pool # |
|---|---|---|---|---|---|
| S1 | GYLQPRTFL | 9 | 268 | 276 | 4 |
| S2 | AYSNNSIAI | 9 | 706 | 714 | 10 |
| S3 | IYQAGSTPCNGV | 12 | 472 | 483 | 5 |
| S4 | FTISVTTEI | 9 | 718 | 726 | 10 |
| S5 | IYQTSNFRV | 9 | 312 | 320 | 10 |
| S6 | IYQAGSTPC | 9 | 472 | 480 | 5 |
| S7 | QYIKWPWYI | 9 | 1208 | 1216 | 6 |
| S8 | CYGVSPTKL | 9 | 379 | 387 | 11 |
| S9 | PPIKDFGGFNF | 11 | 792 | 802 | 11 |
| S10 | VGYQPYRVVVL | 11 | 503 | 513 | 7 |
| S11 | KYNENGTIT | 9 | 278 | 286 | 4 |
| S12 | GYQPYRVVV | 9 | 504 | 512 | 7 |
| S13 | QYGSFCTQL | 9 | 755 | 763 | 8 |
| S14 | SYQTQTNSP | 9 | 673 | 681 | 8 |
| S15 | YQPYRVVVL | 9 | 505 | 513 | 7 |
| S16 | WPWYIWLGF | 9 | 1212 | 1220 | 6 |
| S17 | VYAWNRKRI | 9 | 350 | 358 | 9 |
| S18 | CGPKKSTNL | 9 | 525 | 533 | 9 |
| S19 | KYFKNHTSP | 9 | 1154 | 1162 | 9 |

### Table S2

Table S2. Selected SARS-CoV-2-S peptides with predicted MHC class II (IAd and IEd) restriction

| Peptide ID | Peptide | Length | Start | End | Pool # |
|---|---|---|---|---|---|
| S20 | TRFASVYAWNRKRIS | 15 | 345 | 359 | 1 |
| S21 | RFASVYAWNRKRISN | 15 | 346 | 360 | 1 |
| S22 | FASVYAWNRKRISNC | 15 | 347 | 361 | 1 |
| S23 | INITRFQTLLALHRS | 15 | 233 | 247 | 2 |
| S24 | NYLYRLFRKSNLKPF | 15 | 450 | 464 | 2 |
| S25 | LIRAAEIRASANLAA | 15 | 1012 | 1026 | 3 |
| S26 | NYNYLYRLFRKSNLK | 15 | 448 | 462 | 2 |
| S27 | ASVYAWNRKRISNCV | 15 | 348 | 362 | 1 |
| S28 | IRAAEIRASANLAAT | 15 | 1013 | 1027 | 3 |
| S29 | GNYNYLYRLFRKSNL | 15 | 447 | 461 | 2 |
| S30 | AAEIRASANLAATKM | 15 | 1015 | 1029 | 3 |
| S31 | GGNYNYLYRLFRKSN | 15 | 446 | 460 | 2 |
| S32 | RAAEIRASANLAATK | 15 | 1014 | 1028 | 3 |
| S33 | ATRFASVYAWNRKRI | 15 | 344 | 358 | 1 |
| S34 | NATRFASVYAWNRKR | 15 | 343 | 357 | 1 |

### Fig. S3

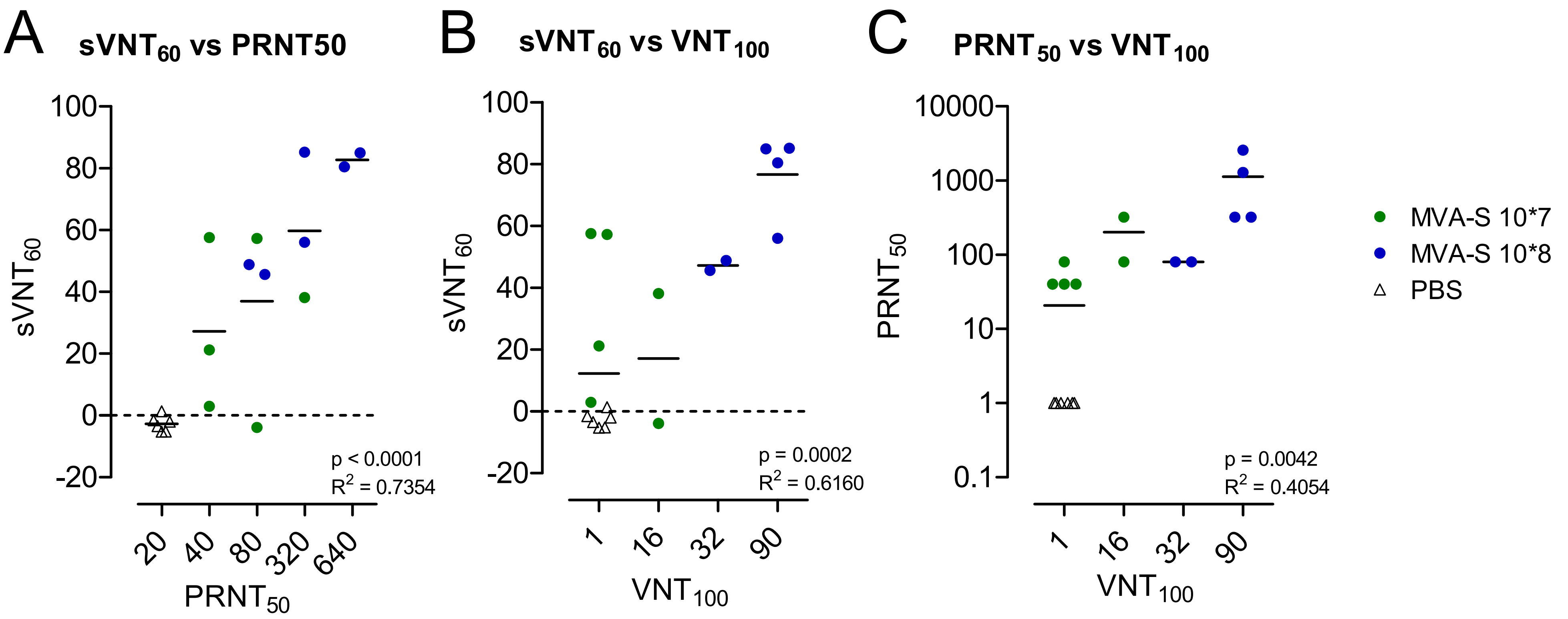


Fig. S3. Robust positive correlation of SARS-CoV-2 neutralizing antibody levels measured by $PRNT_{50}$, $VNT_{100}$ and $sVNT_{1:60}$. Correlation between **(A)** the percent inhibition of SARS-CoV-2 S RBD binding to ACE2 at a 1:60 serum dilution ($sVNT_{60}$) and $PRNT_{50}$ titers, **(B)** $sVNT_{60}$ and $VNT_{100}$ titers; and **(C)** $PRNT_{50}$ and $VNT_{100}$ titers. Correlation was done with one-way ANOVA followed by a test for the trend.

### Fig. S4

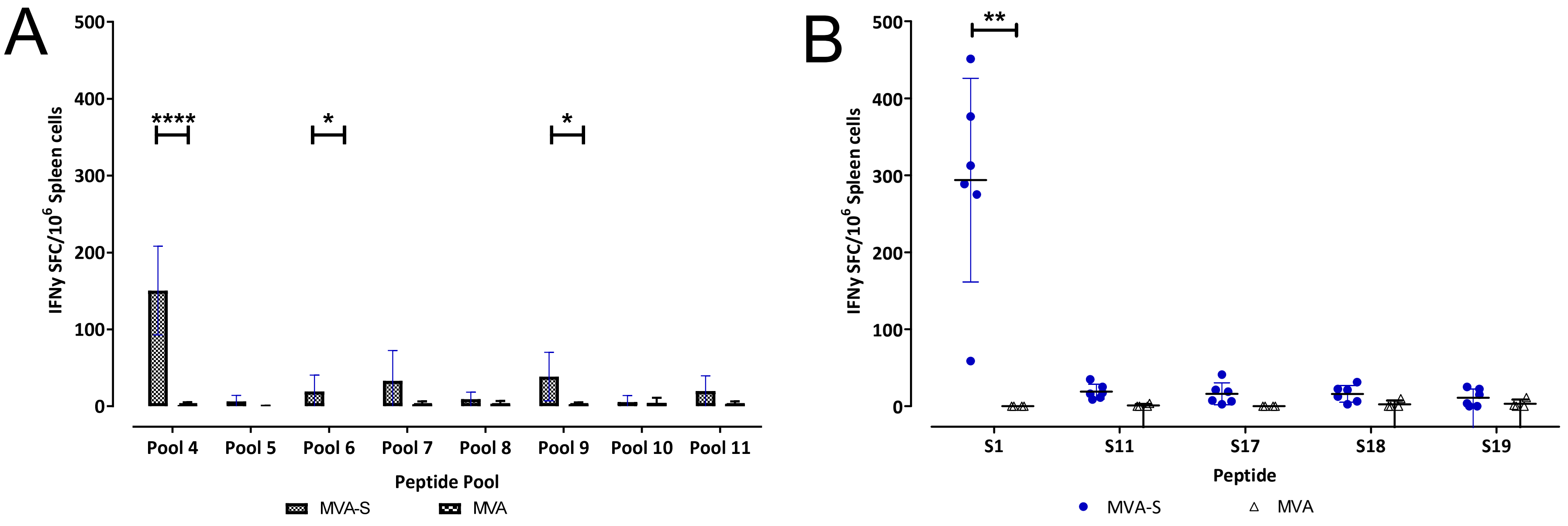


Fig. S4. Identification of H2-d restricted T cell epitopes of SARS-2-S protein. Groups of BALB/c mice (n = 4 to 6) were immunized once with 10*8 PFU MVA-SARS-2-S (MVA-S) or non-recombinant MVA (MVA) via the intramuscular (i.m.) route. Splenocytes were collected and prepared 8 days after the immunization. Total splenocytes were stimulated with pools of 9-12mer peptides or individual peptides from positive pools and were measured by IFN-y ELISPOT assays. **(A)** IFN-y spot forming colonies (SFC) for stimulated splenocytes measured by ELISPOT assays after stimulation with peptide pools (2 to 3 peptides/pool). **(B)** IFN-y SFC for stimulated splenocytes measured by ELISPOT assays after stimulation with each individual peptide from positive pools P4 and P9. Differences between MVA-S and MVA groups per peptide or peptide pool were analyzed by unpaired two-tailed t tests. Asterisks represent statistically significant differences between two groups: * $p < 0.05$; ** $p < 0.01$; **** $p < 0.0001$.

### Fig. S5

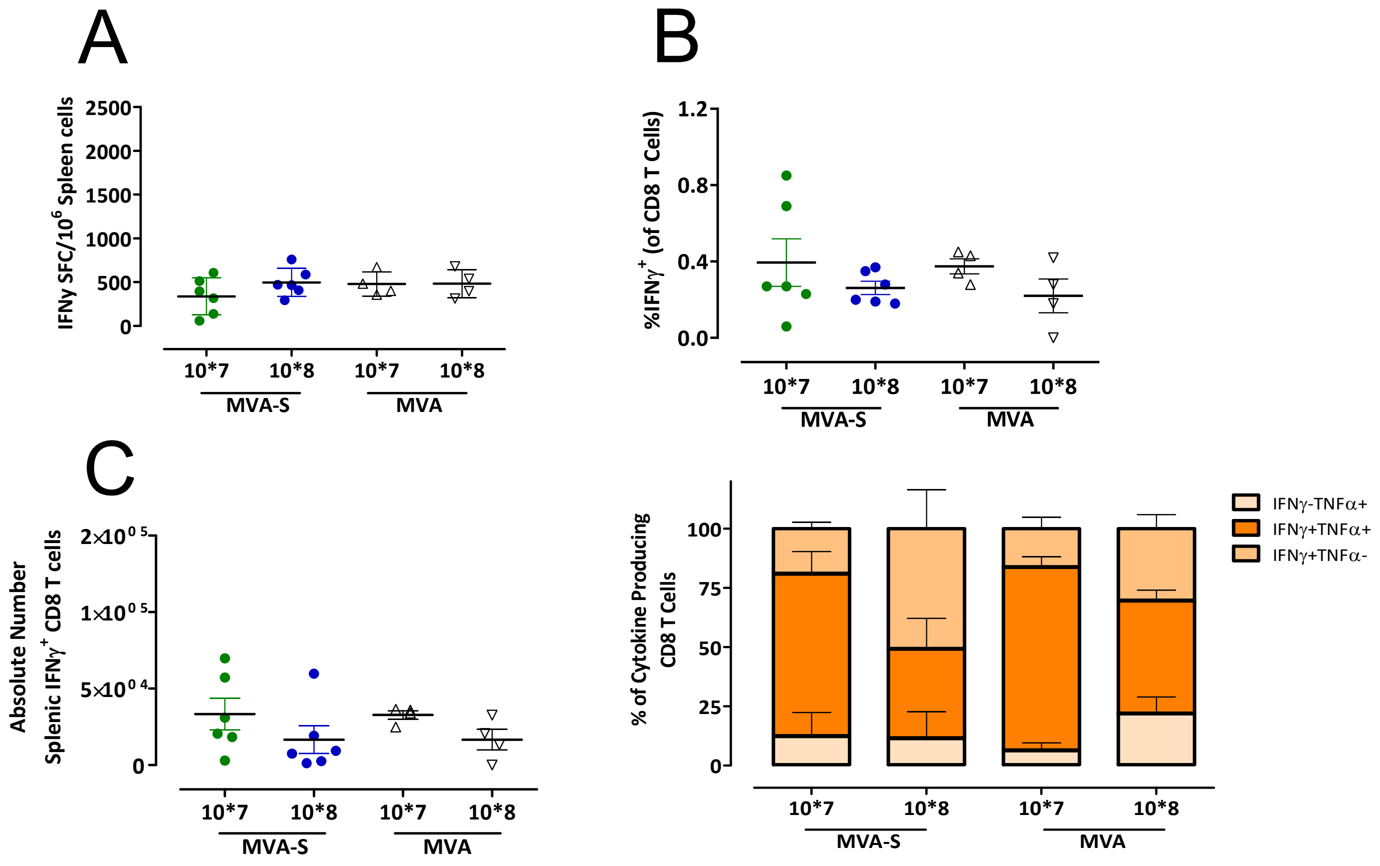


Fig. S5. Activation of MVA-specific CD8+ T cells after prime immunization with MVA-SARS-2-S. Groups of BALB/c mice (n = 4 to 6) were immunized once with 10*7 (LD) or 10*8 (HD) PFU MVA-SARS-2-S (MVA-S) or non-recombinant MVA (MVA) via the i.m. route. Splenocytes were collected and prepared 8 days after the immunization. Total splenocytes were stimulated with the H2d restricted MVA-specific peptide F2(G)$_{26-34}$ and measured by IFN-γ ELISPOT assays and IFN-γ and TNF-α ICS plus FACS analysis. **(A)** IFN-γ spot forming colonies (SFC) for stimulated splenocytes measured by an ELISPOT assay. **(B)** IFN-γ production by CD8+ T cells measured by FACS analysis. Graphs show the frequency and absolute number of IFN-γ+ CD8+ T cells. **(C)** Cytokine profile of F2(G)$_{26-34}$ -specific CD8+ T cells. Graphs show the mean frequency of IFN-γ-TNF-α+, IFN-γ+TNF-α+ and IFN-γ+TNF-α- cells within the cytokine positive CD8 T cell compartment.

### Fig. S6

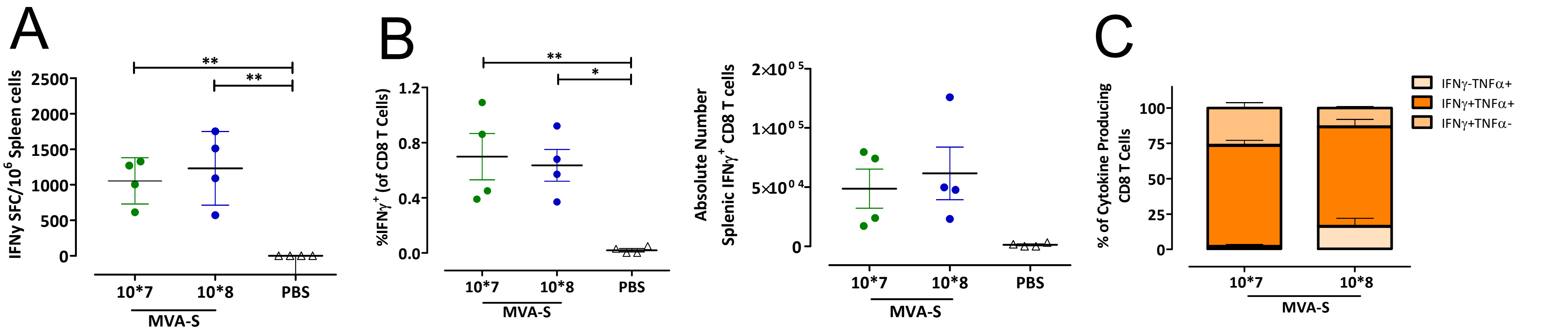


Fig. S6. Activation of MVA-specific CD8+ T cells after prime-boost immunization (21-day interval) with MVA-SARS-2-S. Groups of BALB/c mice (n = 4) were immunized twice with 10*7 (LD) or 10*8 (HD) PFU MVA-SARS-2-S (MVA-S) or non-recombinant MVA (MVA) over a 21-day interval via the i.m. route. Splenocytes were collected and prepared 8 days after the final immunization. Total splenocytes were stimulated with the H2d restricted MVA-specific peptide $F2(G)_{26\text{-}34}$ and were measured by IFN-γ ELISPOT assays and IFN-γ and TNF-α ICS plus FACS analysis. **(A)** IFN-γ SFC for stimulated splenocytes measured by ELISPOT assay. **(B)** IFN-γ production by CD8+ T cells measured by FACS analysis. Graphs show the frequency and absolute number of IFN-γ+ CD8+ T cells. **(C)** Cytokine profile of $F2(G)_{26\text{-}34}$ -specific CD8+ T cells. Graphs show the mean frequency of IFN-γ-TNF-α+, IFN-γ+TNF-α+ and IFN-γ+TNF-α- cells within the cytokine positive CD8+ T cell compartment. Differences between groups were analyzed by one-way ANOVA and Tukey post-hoc tests. Asterisks represent statistically significant differences between two groups: * $p < 0.05$; ** $p < 0.01$.

### Fig. S7

At day 8 after the last immunization we analyzed responses in prime-boost vaccinated animals using three pools (Pool 1-3) of peptides with predicted capacity to activate CD4+ T cells in BALB/c mice. Following restimulation of splenocytes and counting IFN-γ SFC by ELISPOT we detected elevated numbers of IFN-γ-producing cells in splenocytes from MVA-SARS-2-S immunized animals compared to controls.

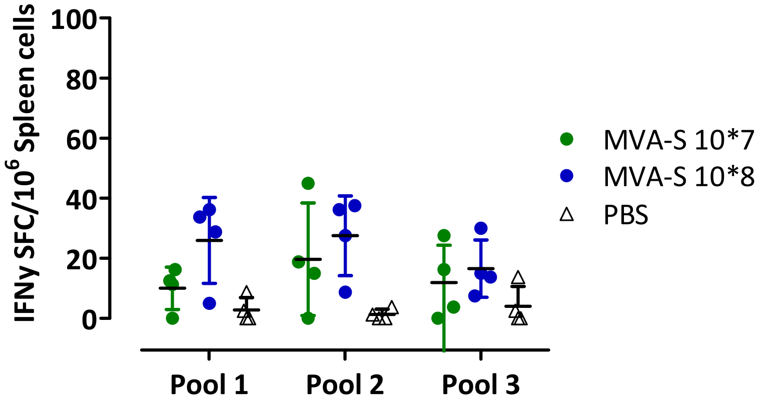


Fig. S7. Testing peptides with predicted capacity to activate CD4+ T cells in BALB/c mice. Groups of BALB/c mice (n=4 to 6) were i.m. immunized in a prime-boost regime (21-day interval) with 10*8 and 10*7 PFU of MVA-SARS-2-S (MVA-S). Mice vaccinated with saline (PBS) were used as a controls. Splenocytes were collected at day 8 post 2nd immunization and stimulated with pools (4 to 6 peptides /pool) of 15mer SARS-2-S derived peptides (Table S2). IFN-γ spot-forming cells were counted by ELISPOT.
